# Ctcf deficiency in myofibers induces pathological genome reprogramming toward the spontaneous development of myopathy

**DOI:** 10.64898/2026.09.28.754564

**Authors:** Jimmy Massenet, Chiara Nicoletti, Luca Caputo, Sören S. Hüttner, Monica Nicolau, Jesus R. Barajas, Maira Rossi, Yuqing Yang, Sara Ancel, Yu Xin Wang, Edoardo Malfatti, Tom H. Cheung, Michael Witcher, Colin Crist, Pier Lorenzo Puri

## Abstract

How perennial, postmitotic multinucleated tissues, such as skeletal myofibers, maintain their identity and transcriptional adaptation to homeostatic perturbations through adult life is an outstanding question. To address this issue, we investigated the consequences of loss of 3D-genome architecture in skeletal muscles by generating myofiber-specific *Ctcf-*deficient (*Ctcf*^mKO^) mice. *Ctcf*^mKO^ mice did not exhibit muscular phenotype at birth but spontaneously developed a severe myopathy. Integrated analysis of snRNAseq, ATACseq and promoter-capture Hi-C revealed both common and fiber-type specific patterns of dysregulated gene expression associated with alterations in chromatin accessibility and promoter-based interactions in Ctcf-deficient myonuclei at distinct stages of myopathy development. Decreased chromatin accessibility at promoters and changes in their connectivity with distal elements were observed across all myonuclei as a direct consequence of *Ctcf* deficiency at early stages and associated with downregulation of genes implicated in myofiber contraction and anabolism, metabolism, adhesion and neuromuscular transmission. Conversely, at later stages, upregulation of genes leading to persistent activation of ER stress/UPR and catabolism resulted from global reconfiguration of chromatin structure and connectivity, partly as indirect consequence of *Ctcf* deficiency. Notably, type-IIB myonuclei exhibited specific alterations in gene expression that culminated in loss of fiber-type identity and ectopic expression of inflammatory genes. These results reveal a requirement of Ctcf for maintenance of fiber-type identity and transcriptional adaptation *in vivo*, through multilayered control of 3D genome integrity. They also indicate an unprecedented association between *Ctcf* deficiency in myofibers and susceptibility to develop myopathies, whereby *Ctcf* dispensability for developmental myogenesis confers vulnerability to develop myopathic syndromes.

## Introduction

Maintenance of fitness and performance in highly dynamic tissues and organs, such as liver, skin, intestine or bone marrow, through the lifespan often relies on their cellular turnover and/or intrinsic regenerative properties(*1*, *2*). In contrast, perennial tissues, such as skeletal muscle, heart and brain, are composed of terminally differentiated, post-mitotic cells that possess very limited cellular turnover and no intrinsic regeneration ability(*3*). It is unknown how these tissues maintain their fitness and performance as well as the ability to adapt to homeostatic perturbations throughout the lifespan.

Skeletal myofibers represent a prototype of perennial terminally differentiated, postmitotic contractile units of skeletal muscles that exist throughout the lifespan. Preserving healthy skeletal muscle mass and performance relies on the balance between catabolic and anabolic pathways that regulate myofiber size as well as on the activation or repression of functional networks that allow them to adapt their performance in response to daily homeostatic perturbations, such as contractile, metabolic and hormonal stimuli, among others(*4–6*). Although skeletal myofibers have no intrinsic regenerative potential, their extrinsic regeneration ability (*e.g.*, in response to injuries or myotrauma) is warranted by a population of muscle stem cells (MuSCs) – also known as satellite cells(*7*, *8*). A peculiar feature of skeletal myofibers is their multinucleation, as each myofiber is composed by hundreds of nuclei endowed with distinct specialization depending on their anatomical position(*9–12*). However, these myonuclei are irreversibly withdrawn from the cell cycle, which accounts for the very limited nuclear turnover potential of adult myofibers, at least in conditions of homeostasis. Thus, a key aspect in preserving skeletal muscle fitness and function through the lifespan relates to the ability of “perennial, postmitotic myonuclei” to warrant long-term maintenance of transcriptional fidelity and competence of myofibers to adapt their transcriptional output upon exposure to homeostatic perturbations.

Studies in the last decade have revealed a novel layer of complexity in the control of gene expression, by identifying essential features of 3D genome architecture. In particular, the recent advent of chromosome conformation capture (3C)-based technologies enabled the analysis of high-order chromatin interactions that revealed essential features of the 3D genome architecture(*13–16*). The 3D genomic architecture is established by high-order chromatin interactions that mediate contacts between functional and structural elements of the genome and promote formation of topological domains for spatial control of gene expression. Briefly, interactions between functional elements - *i.e.*, promoter/enhancer (P/E) interactions – are typically constrained within nuclear domains, such as topological associating domains (TADs as well as sub-TADs called Insulated Neighborhoods - INs), which are defined by high-order chromatin contacts(*17*, *18*). The importance of maintaining the integrity of 3D genome architecture for proper gene expression in different cell types is well illustrated by the growing number of human diseases caused by developmental or post-natal alterations in 3D genome organization(*19*, *20*). Thus, an outstanding topic is to understand how the integrity of 3D genome architecture in postmitotic myonuclei is maintained to support skeletal myofibers fitness and performance from embryonic development through adult life. Specifically, how 3D genome architecture supports myofibers ability to adapt/respond to homeostatic perturbations – i.e., changes in contractile or metabolic demands – by adapting their transcriptional output, remains a poorly investigated topic.

CTCF is a DNA binding protein which regulates, together with the Cohesin complex, the formation of chromatin interactions that define boundaries of nuclear domains, such as TADs and sub-TADs, as well as contacts/loops between cis-regulatory elements of the genome (P/E) that regulate activation, duration and magnitude of gene expression(*21–23*). Most of the studies on CTCF-mediated regulation of 3D genome architecture have been performed in proliferating, undifferentiated cells, especially tissue progenitor cells and cancer cells, or during stem cell differentiation into specific lineages and in models of somatic cell trans-differentiation(*24*, *25*).

Interestingly, while systemic genetic deficiency of CTCF is not tolerated and causes embryonic lethality in mice, CTCF haploinsufficiency in humans has been associated to developmental syndromes, collectively defined to as CTCF-related disorders(*26–28*), which often include musculoskeletal defects and associated symptoms, as they were annotated in over 50% of the patients analyzed(*28*). CTCF has been implicated in the regulation of skeletal myogenesis from myogenic progenitors(*29–33*); however, whether, how and to what extent CTCF contributes to the maintenance of the 3D genome architecture and optimal transcriptional output in post-mitotic skeletal myofibers, through the lifespan is currently unknown.

Here we have addressed this issue by generating a mouse model of Ctcf deficiency in myofibers, which revealed an essential role of Ctcf in the maintenance of 3D chromatin organization and specific gene expression networks, whose dysregulation leads to the spontaneous development of a severe myopathy.

## Results

To investigate the effect of loss of integrity of the 3D genome organization in skeletal myofibers *in vivo*, we exploited a mouse model of myofiber-specific constitutive deficiency of *Ctcf*, in which genetic ablation of endogenous *Ctcf* was induced by Cre-driven deletion of exon 8. To this purpose, we crossed transgenic mice expressing the *Cre* recombinase gene driven by the human alpha-skeletal actin (HSA or *ACTA1*) promoter (HSA^Cre/+^; *Ctcf*^fl/+^ mice) with *Ctcf*^fl/+^ mice containing a *loxP*-flanked sequence at *Ctcf* locus. The offsprings HSA^Cre/+^; *Ctcf*^fl/fl^ mice, herein referred to as *Ctcf*^mKO^ mice (Fig. S1. a) showed efficient Cre-mediated recombination that led to drastic reduction of *Ctcf* transcripts (Fig. S1. b) and CTCF protein (Fig. S1. c) in adult skeletal muscles. As control, CTCF was expressed in the liver of *Ctcf*^mKO^ mice (Fig. S1. c). To ascertain that *Ctcf* deficiency was specifically induced in the nuclei of myofibers (defined as myonuclei) of *Ctcf*^mKO^ muscles, we performed an immunofluorescence analysis, which illustrates how CTCF protein could be detected in nuclei of cell types located within the interstitial space between myofibers or below the basal lamina, corresponding to the location of the muscle satellite cells, but not in myonuclei of *tibialis anterior* (*TA*) muscles of *Ctcf*^mKO^ mice, as defined by the expression of PCM1(*34*) (Fig. S1d). At variance with systemic deficiency of *Ctcf*, which leads to developmental lethality(*35*), *Ctcf*^mKO^ mice were born at the expected mendelian ratio, were viable, and did not manifest any apparent phenotype at birth; however, they exhibited a progressive and spontaneous reduction in body size (Fig. 1a) and weight (Fig. 1b) starting after week 7 of age and invariably died between weeks 14 and 15 of age (Fig. S1. e). As *Ctcf* deficiency was restricted to skeletal myofibers, and because skeletal muscles account for the largest amount of body mass in mammals, we reasoned that the progressive reduction in body size and weight could be caused by loss of muscle mass. Indeed, skeletal muscles (*e.g*. the *TA* and the *gastrocnemius* - *GA*) of *Ctcf*^mKO^ mice exhibited a spontaneous and progressive reduction in the size (Fig. 1c) and weight (Fig. 1d) between 7 and 13 weeks of age, as compared to HSA^cre^ control mice (Ctrl). At the histological level, a proportional reduction of cross-sectional area (CSA) of the myofibers was observed in skeletal muscles of *Ctcf*^mKO^ mice, as compared to controls (Fig. 1e-g). Other skeletal muscles, such as diaphragm, also showed a similar atrophic phenotype (data not shown). Consistently, *Ctcf*^mKO^ mice experienced a progressive loss in exercise endurance ability (Fig. 2a), and by 13 weeks of age their muscles exhibited drastic reduction in peak tetanic force (Fig. 2b) and specific tetanic force (Fig. 2c). Immunofluorescence analysis revealed that these changes in anatomical, morphological and functional parameters of *Ctcf*^mKO^ muscles coincided with a progressive loss of type IIA and IIB fiber and a consensual formation of fibers that did not express any known fiber type-specific myosin and were therefore indicated as “other fibers” (Fig. 2d-e). Furthermore, progressive loss of neuromuscular junctions (NMJs) morphological integrity was observed in muscles of *Ctcf*^mKO^ mice at 7 and 13 weeks of age, as compared to Ctrl mice (Fig. 2f-g).

**Fig. 1:**
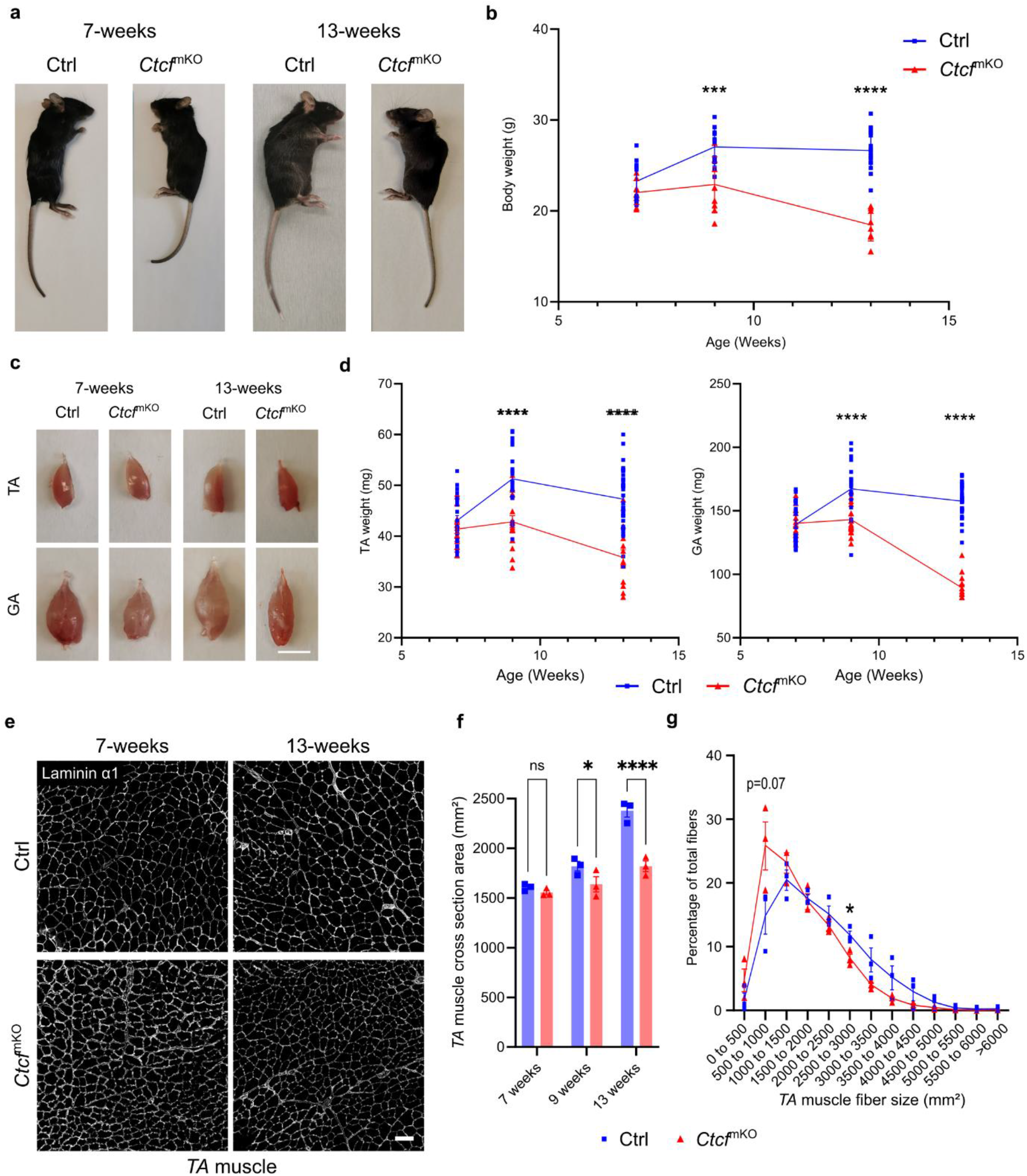
*Ctcf* depletion in myofibers results in progressive atrophy. **a**, Representative images of Ctrl and *Ctcf*^mKO^ mice at 7 and 13 weeks. **b**, Body weight of Ctrl (blue squares) and *Ctcf*^mKO^ (red triangles) mice at 7, 9 and 13 weeks. *n* = 8-15 mice; bars are mean ±SEM; *P* values from two-tailed unpaired T test. **c**, Representative images of the *Tibialis anterior* (*TA*) muscles (top) and *Gastrocnemius* (*GA*) muscles (bottom) of Ctrl and *Ctcf*^mKO^ mice at 7 and 13 weeks; scale bar = 5mm. **d**, Weight of the *TA* muscles (left) and *GA* muscles (right) of Ctrl (blue squares) and *Ctcf*^mKO^ (red triangles) mice at 7, 9 and 13 weeks. *n* = 15-25 muscles; bars are mean±SEM; *P* values from two-tailed unpaired T test. **e**, Representative immunostaining of Laminin α1 in transversal section of the *TA* muscle of Ctrl and *Ctcf*^mKO^ at 7 (left) and 13 weeks (right); scale bar = 100µm. **f**, Cross-sectional area (CSA) of *TA* muscle fibers in Ctrl (blue squares) and *Ctcf*^mKO^ (red triangles) at 7, 9 and 13 weeks. *n* = 3 mice; bars are mean±SEM; *P* values from 2ways-ANOVA with Tukey’s multiple-comparisons test. **g**, Repartition of *TA* fibers CSA in Ctrl (blue squares) and *Ctcf*^mKO^ (red triangles) at 13 weeks. *n* = 3 mice; bars are mean±SEM; *P* values from unpaired T test. \**P*<0.05; *** *P*<0.005; **** *P*<0.001; ns = non significant.

**Fig. 2:**
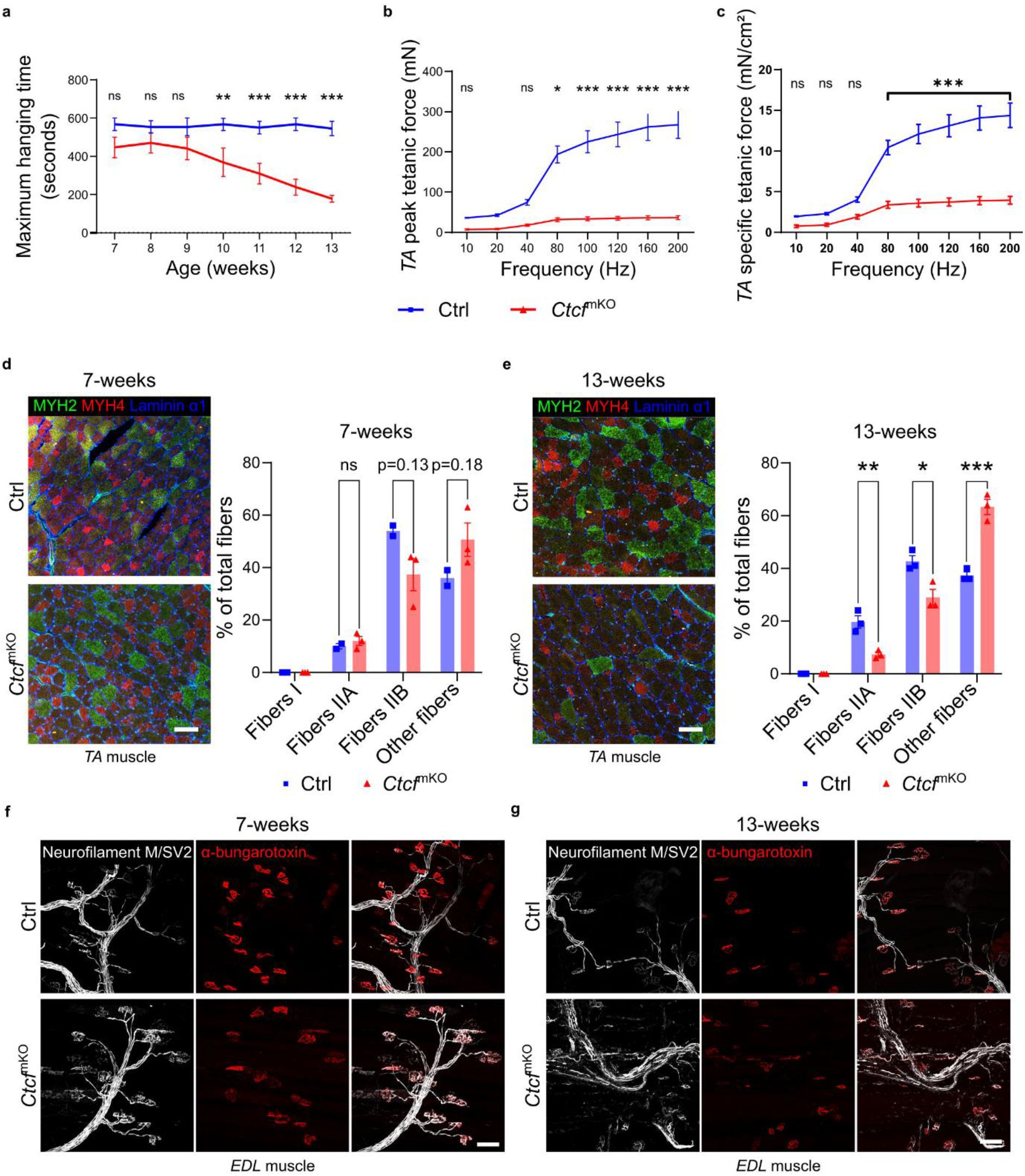
*Ctcf* depletion reduces muscle strength by altering fiber type and neuromuscular junction structure. **a**, Maximum hanging time of Ctrl (blue) and *Ctcf*^mKO^ (red) mice from 7 to 13 weeks. *N* = 8; bars are mean±SEM; *P* values of unpaired T test. **b**, Peak tetanic force of Ctrl (blue) and *Ctcf*^mKO^ (red) mice *Tibialis anterior (TA)* muscle at 13 weeks. *n* = 3; bars are mean±SEM; *P* values from two-tailed unpaired T test. **c**, Specific tetanic force normalized to *TA* weight of Ctrl (blue) and *Ctcf*^mKO^ (red) mice at 13 weeks. *n* = 3; bars are mean±SEM; *P* values from two-tailed unpaired T test. **d**, On the left, representative images of immunostaining of transversal section of *TA* muscle of Ctrl and *Ctcf*^mKO^ at 7 weeks, MYH2 (fiber type **II**A) in green, MYH4 (fiber type IIB) in red and Laminin α1 in blue. On the right, quantification of the percentage of the different fiber types in Ctrl (blue squares) and *Ctcf*^mKO^ (red triangles) at 7 weeks; when no fiber type could have been identified, labeled as “other fibers”. *n* = 2-3 mice; bars are mean±SEM; *P* values from two-tailed unpaired T test; scale bar = 100µm. **e**, On the left, representative images of immunostaining of transversal section of *TA* muscle of Ctrl and *Ctcf*^mKO^ at 13 weeks, MYH2 (fiber type **II**A) in green, MYH4 (fiber type IIB) in red and Laminin α1 in blue. On the right, quantification of the percentage of the different fiber types in Ctrl (blue squares) and *Ctcf*^mKO^ (red triangles) at 13 weeks; when no fiber type could have been identified, labeled as “other fibers”. *n* = 3 mice; bars are mean±SEM; *P* values from two-tailed unpaired T test; scale bar = 100µm. **f-g**, Representative images of immunostaining of isolated myofiber bundles of *EDL* muscle of Ctrl and *Ctcf*^mKO^ at 7 weeks (**f**) and 13 weeks (**g**), with α-bungarotoxin (nicotinic acetylcholine receptors) in red, Neurofilament-M and synaptic vesicle glycoprotein 2A (SV2A) in white. scale bar = 100µm. \**P*<0.05; ** *P*<0.01; *** *P*<0.005; ns = non significant.

Because the Cre recombinase is driven by the HSA promoter, which induces gene expression by 9.5 days *post coitum* (*p*.*c*.)(*36*), a stage in which embryonic myogenic progenitors begin to form primary myofibers(*37*), we considered the possibility that the severe skeletal muscle phenotype observed in *Ctcf*^mKO^ mice during postnatal life could arise from an impaired developmental myogenesis caused by *Ctcf* deficiency. Thus, although *Ctcf*^mKO^ homozygous mice were born with apparently normal musculature, we investigated whether a latent or subtle impairment of developmental myogenesis could determine a delayed or defective postnatal growth of skeletal muscles. However, a comparative analysis of embryonic skeletal muscle development did not reveal any noticeable histological difference between *Ctcf*^mKO^ and Ctrl control embryos, as they showed similar patterns of MyHC expression, distribution and cross-sectional area, as well as cell proliferation (as judged by the frequency of Ki67 positive nuclei within muscles) and apoptosis (as judged by the frequency of cleaved caspase 3 positive cells within muscles) at day 14.5 *p*.*c*. (Fig. S2. a-e) or day 16.5 *p*.*c*. (Fig. S2. f-j). To conclusively rule out the potential impact of a latent developmental phenotype caused by *Ctcf* deficiency, we sought to compare the effect of constitutive *Ctcf* deficiency, as driven by developmental activation of HSACre in *Ctcf*^mKO^, with conditional induction of *Ctcf* deficiency during post-natal life. To this purpose, conditional Ctcf^fl/fl^ mice were crossed with tamoxifen (TAM) inducible skeletal muscle specific Cre recombinase (HSA^merCREmer/merCREmer^) mice to generate a tamoxifen (TAM)-inducible *Ctcf*^mKO^ mouse line (*Ctcf*^mKO^ TAM) and relative HSA^merCREmer/merCREmer^ control (Ctrl TAM). Fig. S3 shows that *Ctcf* deficiency was effectively induced by TAM administration to 4-month-old *Ctcf*^mKO^ TAM mice, leading to a drastic reduction in CTCF levels by 2 weeks and 4 months after TAM induction, as compared to Ctrl TAM (Fig. S3. a) and resulting in about 75% decrease in *Ctcf* transcripts downstream of exon 8 at both time points (Fig. S3. b-c). *Ctcf*^mKO^ TAM mice exhibited a phenotype characterized by a progressive reduction in body (Fig. S3. d) and muscle (Fig. S3. e-f) weight, with development of myofiber atrophy (Fig. S3. g), upon TAM treatment, similar to *Ctcf*^mKO^ mice (shown in Fig. 1). We then compared the changes in gene expression induced by constitutive or conditional deficiency of *Ctcf*, by performing bulk RNAseq analysis of *GA* muscles from *Ctcf*^mKO^ or *Ctcf*^mKO^ TAM mice, as compared to their respective controls. Principal component analysis (PCA) shows that, beyond the batch effect differences between the two mouse lines, as accounted by PC1 (data not shown), PC2 could discriminate samples from control mice, which all clustered together, from samples of *Ctcf*^mKO^ mice, which positioned distant from controls, according to the timing of *Ctcf* deficiency (Fig. S4. a). Specifically, 7 weeks *Ctcf*^mKO^ samples positioned closer to controls, while 13 weeks *Ctcf*^mKO^ samples and 4 months treated *Ctcf*^mKO^ TAM samples clustered more distant (Fig. S4. a), indicating similar changes in gene expression profile induced by long-term constitutive or conditional *Ctcf* deficiency. Consistently, gene ontology (GO) analysis revealed analogies in the biological process related to up- and down-regulated genes from samples of *Ctcf*^mKO^ or *Ctcf*^mKO^ TAM mice. Indeed, upregulation of genes implicated in endoplasmic reticulum (ER) stress/unfolded protein response (UPR), negative regulation of gene expression, immune response/inflammation and apoptosis, and downregulation of genes implicated in muscle contraction/development, glucose metabolism and mitochondrial activity, were detected both in samples from 4 months treated *Ctcf*^mKO^ TAM (Fig. S4. b) and *Ctcf*^mKO^ mice along the transition from 7 to 13 week of age (Fig. S4. c-d). Thus, conditional induction of *Ctcf* deficiency in post-natal life could replicate salient morphological and transcriptional features of constitutive *Ctcf* deficiency. Interestingly, different from *Ctcf*^mKO^ mice, which invariably died between week 14 and 15 of age, *Ctcf*^mKO^ TAM could survive beyond 13 weeks following TAM-induced *Ctcf* deficiency.

Overall, the bulk RNA-seq from whole muscles indicates that myofiber-specific *Ctcf* deficiency leads to extensive alterations in gene expression, consisting of progressive loss of contractile and metabolic genes and parallel activation of stress response and catabolic processes that converge into the development of the final phenotype of myofiber atrophy. It also reveals changes in the expression of inflammatory genes, which can either derive from a cell autonomous ectopic gene expression from *Ctcf* deficient myofibers or from non-cell autonomous transcriptional alterations in other cell types, such as muscle-resident inflammatory cells.

To further investigate the cell-autonomous effect of CTCF deficiency in myofibers, we focused our analysis on myonuclei. We used PCM1 as a myonuclear specific protein to isolate myonuclei from skeletal muscles(*38*). Given the established role of CTCF in regulating 3D genome architecture and chromatin structure(*22, 39–45*), we first investigated whether *Ctcf* deficiency led to alterations in chromatin interactions and accessibility, by performing promoter capture Hi-C (pcHiC) and ATAC-seq analysis in myonuclei of *GA* muscles isolated from *Ctcf*^mKO^ mice at 7 and 13 weeks of life, as compared to Ctrl mice. PCA from ATAC-seq analysis of myonuclei isolated from *Ctcf*^mKO^ mice (or Ctrl) shows that while chromatin accessibility did not show significant change from 7 and 13 weeks in Ctrl mice, CTCF deficiency resulted in dynamic changes of chromatin accessibility at 7 and 13 weeks in *Ctcf*^mKO^ mice, as evidenced by PC1 and PC2, respectively (Fig. 3a). Specifically, coherent patterns of reduced chromatin accessibility (Fig. 3b - clusters D and E) were detected at gene promoters (Fig. 3c) in myonuclei of both 7- and 13-week-old *Ctcf*^mKO^ mice, as compared to their respective controls. A more dynamic patterns of changes in chromatin accessibility was detected at other elements of the genome in myonuclei of 7- and 13-week-old *Ctcf*^mKO^ mice (Fig. 3b-c), with specific patterns of increased accessibility transiently detected in myonuclei of 7-week-old *Ctcf*^mKO^ mice and chromatin compaction was restored in 13-week-old *Ctcf*^mKO^ mice (Fig. 3b - cluster B), and new peaks of increased accessibility detected in myonuclei of 13-week-old *Ctcf*^mKO^ mice (Fig. 3b - cluster C). A pattern of progressive increase in accessibility was detected in a few genomic regions of myonuclei at 7 and 13 weeks old *Ctcf*^mKO^ mice (Fig. 3b - cluster A). Finally, a general trend of partial recovery in chromatin accessibility was observed in myonuclei of 13 weeks old *Ctcf*^mKO^ mice (Fig. 3b - cluster D). Overall, these data indicate that CTCF deficiency leads to genome-wide dynamic changes in chromatin accessibility that include both stage-specific and progressive alterations at proximal (e.g. promoters) and distal (e.g. enhancers) regulatory elements. Given the well-established role of CTCF as regulator of 3D genome architecture for topological control of P/E communications(*22–24*), we evaluated whether and to what extent CTCF deficiency altered P/E interactions genome-wide, by performing promoter capture Hi-C (pcHi-C) analysis in myonuclei isolated from 7- and 13-week-old *Ctcf*^mKO^ mice or Ctrl mice. This analysis revealed an increased frequency of differential chromatin interactions between gene promoters and other genomic elements in myonuclei of 7-week-old *Ctcf*^mKO^ mice, as compared to controls (Fig. 3d). While the frequency of these changes was reduced in myonuclei of 13-week- old *Ctcf*^mKO^ mice (Fig. 3d and f), at this stage the length of differential chromatin interactions between gene promoters with other promoters and distal elements was significantly increased (Fig. 3e and f). Overall, these data suggest that CTCF deficiency leads to stepwise alterations in 3D nuclear architecture, whereby initial high frequency in changes of promoter interactions with other genomic elements are followed by an increased number of long-range interactions between promoters and more distally located genomic elements, including promoters or others (*i*.*e*., enhancers). These changes are consistent with the potential alterations of CTCF-regulated boundaries of TADs and subTADs in myonuclei of *Ctcf*^mKO^ mice that are predictive of myofiber-specific extensive alterations in gene expression, as also anticipated by bulk RNA-seq data.

**Fig. 3:**
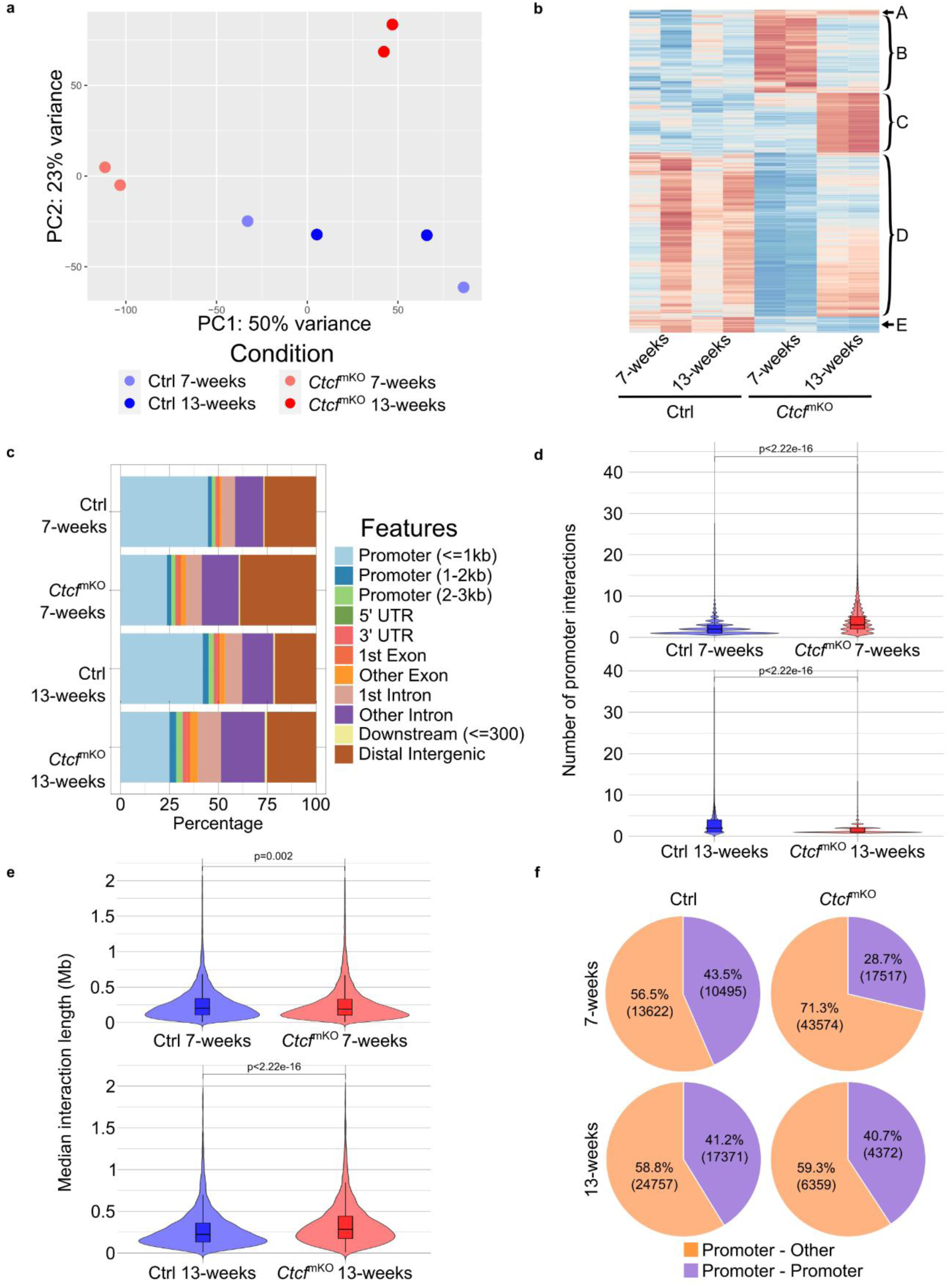
*Ctcf* absence in myofibers causes changes in chromatin accessibility and interactions. **a**, PCA plot of bulk ATAC-seq from isolated myonuclei at 7 and 13 weeks in *Ctcf*^mKO^ and Ctrl. *n* = 2 mice. **b**, Heatmap plot representing differential chromatin accessibility in the comparisons of *Ctcf*^mKO^ versus Ctrl at 7 and 13 weeks. Chromatin accessibility represented as z-scores. *Q* value < 0.05. *n* = 2 mice. **c**, Genome-wide distribution of chromatin accessibility peaks in the different experimental conditions. List of accessibility peaks for each condition were derived through irreproducible discovery rate (IDR) of the biological replicates. *n* = 2 mice. **d**, Violin plots representing changes in number of promoters’ chromatin interaction partners between Ctrl (blue) and *Ctcf*^mKO^ (red), at 7 (top) and 13 weeks (bottom). Statistical significance calculated with Wilcoxon test. *n* = 2 mice. **e**, Violin plots representing changes in chromatin interaction length between Ctrl (blue) and *Ctcf*^mKO^ (red), at 7 (top) and 13 weeks (bottom). Statistical significance calculated with Wilcoxon test. *n* = 2 mice. **f**, Pie chart representing changes in proportion of promoter-promoter (purple) and promoter-other (orange) interactions between Ctrl (left) and *Ctcf*^mKO^ (right), at 7 (top) and 13 (bottom) weeks. *n* = 2 mice.

We then focused on cell nucleus-specific alterations of gene expression by performing single nucleus RNA-seq (snRNA-seq) analysis of muscles isolated from 7- or 13-week-old *Ctcf*^mKO^ or Ctrl mice, to discriminate the cell autonomous effect of CTCF deficiency in myonuclei from non-cell autonomous alterations in nuclei of other muscle-resident cell types. Uniform Manifold Approximation and Projection (UMAP) analysis shows that minimal changes in number of nuclei were observed at 7 weeks in *Ctcf*^mKO^ mice, mostly accounted by a small decrease in number of type IIB myonuclei and a relative increase in FAP number; however, by 13 weeks, myonuclei of *Ctcf*^mKO^ mice exhibited an almost complete loss of type IIB myonuclei, which were replaced by a population of myonuclei that did not display any known “fiber type” identity marker and were indicated as “other myonuclei” (Fig. 4a-b), in analogy to the “other fibers” detected by immunofluorescence in Fig. 2d-e. Of note, at this stage we observed a massive increase in the number of muscle-resident immune cells (Fig. 4a-b), which suggests a spontaneous development of inflammatory myositis, as an indirect consequence of myofiber-specific deficiency in CTCF.

**Fig. 4:**
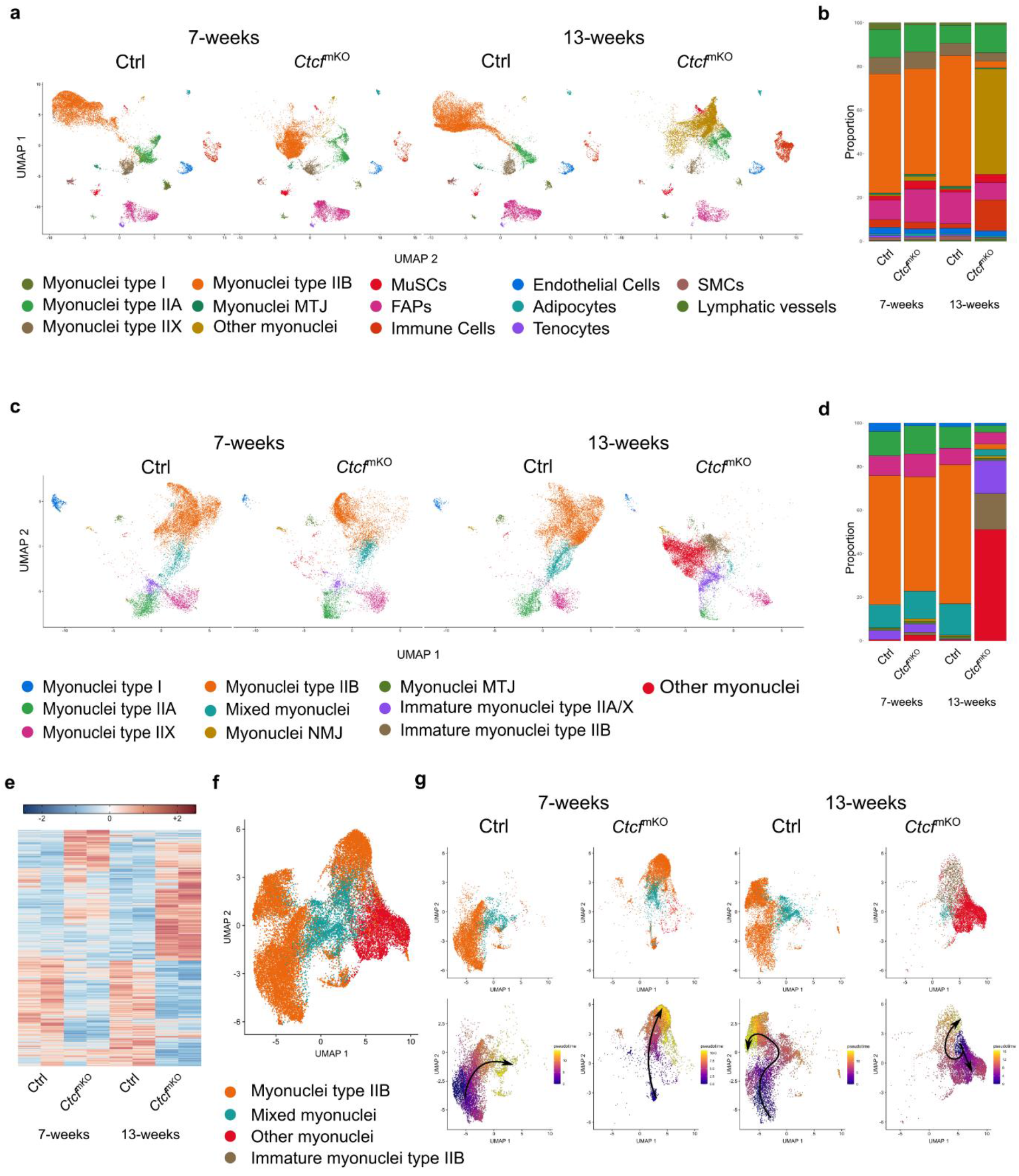
Loss of *Ctcf* in myonuclei drives an alternative transcriptional program. **a**, UMAP embedding of all the snRNAseq conditions: Ctrl and *Ctcf*^mKO^ at 7 and 13 weeks. The dataset is colored and labeled by metacluster. *n* = 2 mice. **b**, Proportion in percentage of the metaclusters in Ctrl and *Ctcf*^mKO^ mice at 7 and 13 weeks. Metacluster labeling as in **a**. *n* = 2 mice. **c**, UMAP of the myonuclei metacluster split by conditions, Ctrl and *Ctcf*^mKO^ at 7 and 13 weeks. The dataset is colored and labeled by myonuclei subclusters. *n* = 2. **d**, Proportion in percentage of the myonuclei subclusters in Ctrl and *Ctcf*^mKO^ mice at 7 and 13 weeks. Subcluster labeling as in **c**. *n* = 2 mice. **e**, Heatmap of differential gene expression generated from the pseudo-bulk analysis of the myonuclei subclusters of Ctrl and *Ctcf*^mKO^ mice at 7 and 13 weeks. *n* = 2 mice. **f**, UMAP of Type IIB, Immature type IIB, Mixed and Other myonuclei clusters. *n* = 2. **g**, On top, UMAP of Type IIB, Immature type IIB, Mixed and Other myonuclei clusters split by conditions (Ctrl and *Ctcf*^mKO^ at 7 and 13 weeks), colored and labeled by subclusters. On the bottom, Monocle3 pseudotime UMAP of Type IIB, Immature type IIB, Mixed and Other myonuclei split by conditions (Ctrl and *Ctcf*^mKO^ at 7 and 13 weeks). *n* = 2 mice.

To identify gene expression changes most directly associated with myofiber-specific CTCF deficiency, we focused our analysis on myonuclei. UMAP analysis of the myonuclei subcluster showed a progressive reduction in type IIB myonuclei (from fast-twitch, anaerobic, glycolytic fibers) of muscles from *Ctcf*^mKO^ mice, which by 13 weeks of age were replaced by the cluster of “other myonuclei” that was not present in Ctrl muscles and accounted for about 50% of the myonuclei in *Ctcf*^mKO^ muscles (Fig. 4c-d). Moreover, muscles of 13-week-old *Ctcf*^mKO^ mice also exhibited a decrease in type IIA myonuclei (e.g. fast-twitch, oxidative-glycolytic, intermediate fibers) as well as myonuclei expressing different marks of fiber types (mixed fibers), with a concomitant increase in number of immature myonuclei that did not express specific markers of type IIA, type IIB or IIX fibers (Fig. 4c-d). The massive changes in myonuclear composition observed in *Ctcf*^mKO^ mice were associated with progressive alterations in gene expression at 7 and 13 weeks of age (Fig. 4e; Fig. S5. a and d) that culminated in a similar number of up- and down-regulated genes in myonuclei of 13-week-old *Ctcf*^mKO^ mice, as compared to their controls (Fig. 4e). Gene ontology analysis revealed that downregulated genes in myonuclei of *Ctcf*^mKO^ mice at both 7 and 13 weeks of age were implicated in processes related to skeletal myogenesis, muscle contraction, sarcomere organization, neuromuscular junctions, glucose uptake and metabolism, as well as glycogen storage and usage (Fig. S5. b and e). Conversely, stepwise upregulation of genes implicated in processes related to ribosomal control of protein synthesis, ER stress/UPR, regulation of cytoskeleton and intracellular signaling, apoptosis and activation of immune response were observed upon CTCF deficiency in myonuclei of *Ctcf*^mKO^ mice at both 7 and 13 weeks (Suppl. Fig.5c and f). Consistent with the patterns of differentially expressed genes (DEGs) from snRNA-seq analysis (Fig. S6.a), western blot analysis revealed increased expression of the phosphorylated form of the main regulator of protein translation EIF2A, as marker of persistent ER stress (Fig. S6. b and c)(*46*, *47*), and elevated levels of UPR effectors(*48*, *49*), such as ATF6 (Fig. S6. b and d), ATF4 (Fig. S6. b and e) and PERK (Fig. S6. b and f) in 13 week old *Ctcf*^mKO^ mice, as compared to controls. Likewise, myofibers of 13-week-old *Ctcf*^mKO^ mice showed increased expression of markers of muscle atrophy, such as 15- PGDH(*50*, *51*) (Fig. S6. b, g and i), downregulation of genes implicated in muscle anabolism, such as BCL6(*52*, *53*) (Fig. S6. b and h) and exhibited massive patterns of ubiquitination (Fig. S6. j). Among the upregulated inflammatory genes, we detected receptors, ligands and intracellular effectors of *Toll like receptors* (*Tlr)*, *Interferon gamma* (*Ifn-g)*, *Tumor Necrosis Factor (Tnf)*-, and *Interleukin (Il)*-regulated immune responses (Fig. S7. a-c).

To investigate the progressive transcriptional shift occurring in Ctcf^mKO^ myonuclei, pseudotime trajectories were inferred for the main myonuclei types of the *GA* muscle; namely, type IIB (Fig 4f-g), type IIA (Fig. S8. a-c), type IIX (Fig. S8. b-d).

Ctrl myonuclei exhibited a steady trajectory at 7 and 13 weeks, whereby a proportion of type IIB fibers progressed toward mixed fibers (Fig. 4g). In contrast, Ctcf^mKO^ myonuclei exhibited severe alterations in pseudotime trajectory, with progressive shift of fiber identity observed at 7 and 13 weeks (Fig. 4g). By 7 weeks myonuclei type IIB transitioned toward a subcluster of type IIB immature myonuclei; while, by 13 weeks, myonuclei exhibited a trajectory that bifurcated into two distinct sub-clusters: immature myonuclei and “other myonuclei” (Fig. 4g). Pseudotime analysis of type IIA (Fig. S8. c) and type IIX (Fig. S8. d) myonuclei from control mice revealed trajectories similar to those observed in type IIB myonuclei, with myonuclei type IIA or IIX progressing toward mixed myonuclei. Conversely, Ctcf^mKO^ myonuclei type IIA (Fig. S8c) and myonuclei type IIX (Fig. S8. d) exhibited an early shift toward mixed myonuclei that could be appreciated by 7 weeks of age and culminated into a massive transition toward immature myonuclei type IIA/IIX at 13 weeks. Overall, these trajectories indicate a myonuclei identity reprogramming underlying the pathological phenotype developed by *Ctcf* deficient muscles.

We therefore narrowed our analysis to the differential patterns of gene expression of pseudobulk myonuclei that underwent the most massive changes from 7 to 13 weeks of *Ctcf*^mKO^ mice - *i*.*e*. myonuclei type IIB (shown in Fig. S9. a and d) and “other myonuclei” (shown in Fig. S11. a and d). Specifically, the cluster of type IIB myonuclei of *Ctcf*^mKO^ mice exhibited a progressive downregulation of genes implicated in skeletal myogenesis, muscle contraction, sarcomere organization, and anabolic response, as well as genes implicated in glucose uptake and metabolism and glycogen storage and usage, between 7 (Fig. S9. b) and 13 (Fig. S9. e) weeks of life. Of note, among the downregulated genes, we annotated genes implicated in muscle development and myogenic identity (such as the *Mef2a, Mef2c, Mef2d, Myf6, Six1* and *Six4*)(*54–57*) (Fig. S10. a-d) and a massive downregulation of genes implicated in the epigenetic activation of transcription muscle-specific genes, including co-activators (such as Set-containing lysine methyltransferases, the *Carm1* arginine methyltransferases, the *Kdm1a/Lsd1* Lysine-specific demethylase 1A, the histone acetyltransferases *Cbp* and *Kat2b-P/Caf*, and the *Mediator Complex Subunit 13l, Med13l*)(*58–60*) and the muscle-specific SWI/SNF complex sub-unit *Smarcd3/Baf60c*(*61*), which also promotes glycolytic muscle metabolism(*62*) (Fig. S10. e). Furthermore, we observed downregulation of receptors and transducers of key signaling, such as *Acvr2a, Bmpr1a, Traf6*, the Spermine oxidase *Smox* (a key enzyme in the polyamine catabolic pathway) and the RNA-induced silencing complex (RISC) component *Ago2* (Fig. S10. e). This extensive downregulation of myogenic determination factors, muscle-specific and general components of the transcriptional machinery, together with receptors and enzymes implicated in key developmental processes, indicates a nuclear reprogramming and rewiring toward the loss of myonuclei type IIB identity. A parallel upregulation of genes implicated in protein translation, endoplasmic reticulum (ER) stress/unfolded protein response (UPR), mitochondrial activity was observed in type IIB fibers of *Ctcf*^mKO^ mice from 7 to 13 weeks (Fig. S9. c and f). By 13 weeks the residual amount of myonuclei type IIB did not cluster, but were rather dispersed (Fig. 4c), according to their more heterogeneous pattern of gene expression suggestive of loss of cell identity. Importantly, these myonuclei showed upregulation of genes implicated in immune response/inflammation (Fig. S9. f) that was not observed in other subclusters of myonuclei, indicating that *Ctcf* deficient myonuclei type IIB are the only source of ectopic activation of inflammatory genes by week 13.

The cluster of “other myonuclei” was represented by a small number of nuclei in muscles of 7-week-old *Ctcf*^mKO^ mice (Fig. 4c and d) that displayed differential gene expression, as compared to controls (Fig. S11. a), and showed downregulation of genes implicated in muscle contraction and anabolic response (Fig. S11. b) and upregulation of genes implicated in various cellular processes (Fig. S11. c). However, by 13 weeks “other myonuclei” accounted for the major population of myonuclei (Fig. 4c and d) and showed a distinctive pattern of gene expression (Fig. S11. d), with further downregulation of genes implicated in myogenic identity (including *Mef2C*, *MyoD1, Myf6 and Smarcd3/Baf60c*) as well as genes implicated in muscle contraction, sarcomere organization, anabolic response, as well as genes implicated in glucose uptake/metabolism and glycogen storage/metabolism (Fig. S11. e). The simultaneous downregulation of these genes likely converges into a global loss of functional and biological properties observed in many myopathies, pointing to *Ctcf* mutations as potential drivers of development and penetrance of muscular disorders. According to the temporal contiguity with type IIB myonuclei loss and/or IIB conversion into “other myonuclei” (*i.e.*, atrophic myonuclei), the atrophic myonuclei exhibit massive upregulation of ER stress/UPR and ensuing processes, including muscle catabolism/atrophy, macro-autophagy and apoptosis (Fig. S11. f).

Interestingly, no ectopic expression of inflammatory genes was observed in atrophic fibers, suggesting that loss of identity of myonuclei type IIB fibers caused by *Ctcf* deficiency leads to transcriptional bifurcation, whereby most of the myonuclei type IIB convert into atrophic myonuclei, with residual type IIB myonuclei that lost their myogenic identity persist and ectopically express pro-inflammatory genes.

We next investigated whether and to what extent these events are directly caused by *Ctcf* deficiency and their potential interdependence at the chromatin level, by an integrated analysis of RNAseq, ATACseq and pc-HiC datasets generated from myonuclei of *Ctcf*^mKO^ mice or controls. To distinguish direct from indirect consequences of *Ctcf* depletion on chromatin organization and transcription, we first characterized the *Ctcf* binding landscape in control myofibers at 7 weeks using myonuclei CTCF ChIP-seq. Integration of these data with chromatin accessibility profiles generated by ATACseq from isolated myonuclei, revealed that genomic regions bound by CTCF in control conditions undergo a marked loss of accessibility upon *Ctcf* deletion, a phenotype already evident at 7 weeks and persisting at 13 weeks (Fig. 5a). This finding indicates that CTCF binding is required to maintain an accessible chromatin state at its target loci and that its loss leads to progressive chromatin compaction. Consistent with this observation, differential accessibility analysis between control and *Ctcf*^mKO^ myonuclei showed an enrichment of CTCF peaks among regions exhibiting reduced accessibility, with a stronger effect observed at promoters, as compared to non-promoter regions (Fig. 5b). This promoter-biased sensitivity suggests a prominent role for CTCF in sustaining promoter accessibility, potentially through insulation, stabilization of local chromatin architecture, or facilitation of transcriptional machinery recruitment(*26*). Indeed, motif analysis of differential ATAC peaks shows only a slight preferential enrichment in binding motifs for transcription factors (TF), such as MEF2 family members and the Serum Response Factor (SRF) accessory protein Elk4, at promoters that show reduced chromatin accessibility in *Ctcf*^mKO^ myonuclei at 7 weeks (Fig. 5c). Although MEF2 and Elk4 can cooperate to activate the expression of muscle genes in myofibers, the lack of enrichment for specific TF-binding motifs at promoters with reduced chromatin indicates that chromatin compaction detected at promoters is a widespread effect of *Ctcf* deficiency on gene promoters, rather than a specific effect caused by loss of *CTCF* cooperation with distinct TFs. In contrast, loss of chromatin accessibility at *CTCF*-bound non-promoter elements, which typically include enhancers, appears driven by myofiber-specific TFs, as it was associated with enrichment of motifs for TFs implicated in the activation of muscle-specific gene expression, such as MEF2 family members, *Six2*, *Bach2* and *Nfil3*, in addition to common enhancer-binding TFs, such as AP1 family members (Fig. 5c). Interestingly, at 13 weeks we observed a partial recovery of chromatin accessibility at non-promoter regions, as also anticipated by ATACseq heatmap shown in Fig. 3b, that was associated with the enrichment of these TFs (Fig. 5c). However, this partial recovery did not reverse the pattern of repression of gene expression.

**Fig. 5:**
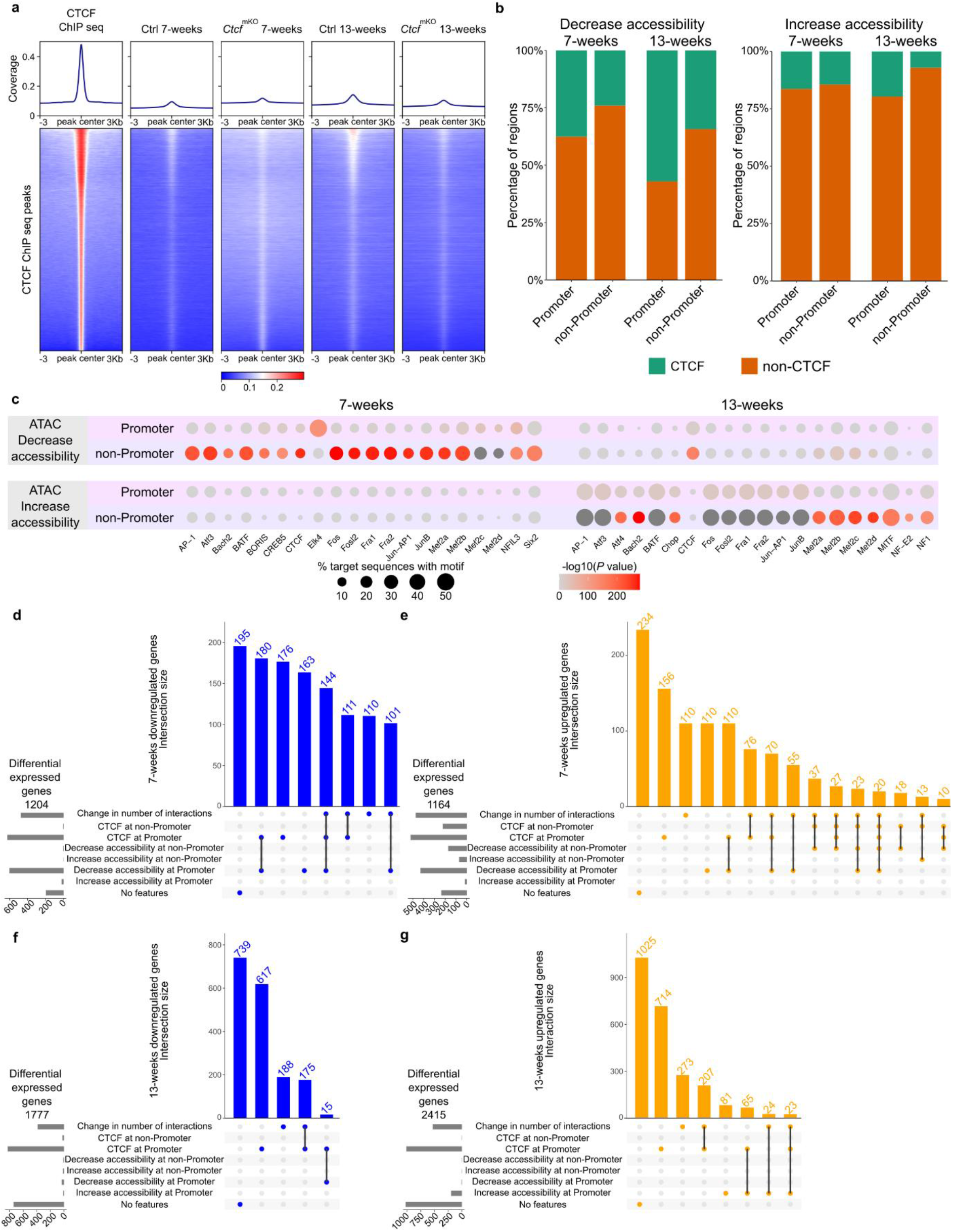
Characterization of direct and indirect transcriptional and epigenetic dysregulation upon *Ctcf* depletion. **a**, Tornado plot of CTCF ChIP-seq signal (left) and chromatin accessibility in Ctrl and *Ctcf*^mKO^ mice at 7 and 13 weeks (right) within CTCF peaks from Ctrl myofibers. *n* = 2 replicates; each replicate is a pool of 2 mice. **b**, Stacked bar plot of the enrichment of CTCF peaks from Ctrl myofibers in differential chromatin accessibility regions (increased accessibility on the left, and decreased accessibility on the right) at 7 and 13 weeks, split by promoter and non-promoter differential regions. **c**, Bubble plot showing the top 20 TFs motifs found enriched within differential chromatin accessibility regions (split in promoter and non-promoter) at 7 and 13 weeks. Bubble size corresponds to % Targets Sequences with Motif, while the color scale represents the statistical significance (-log_10_(p-value)). **d-g**, Upset plots representing the combination of epigenetic dysregulation events connected to differential gene expression, for downregulated gene at 7 weeks (**d**) and 13 weeks (**f**), and upregulated genes at 7 weeks (**e**) and 13 weeks (**g**). Overlap cutoff at 10 events.

To further dissect the regulatory mechanisms underlying the transcriptional changes induced by *Ctcf* loss, we stratified differentially expressed genes (DEGs) according to multiple chromatin features: (i) CTCF binding at gene promoters and/or distal regulatory regions; (ii) changes in chromatin accessibility at promoters and distal elements; and (iii) alterations in chromatin interaction connecting promoters and distal elements, between Ctrl and *Ctcf*^mKO^ conditions. Distal elements were assigned to target promoters using pcHi-C data, enabling an integrated view of 3D genome regulation (Fig. 5d–g). At 7 weeks, downregulated genes showed a strong association with perturbations of chromatin accessibility, with 51% of DEGs directly associated with these perturbations (Fig. 5d). The most prevalent categories included genes with CTCF binding at promoters (15%), loss of promoter accessibility (14%), or both (15%). Importantly, a notable fraction of downregulated genes (12%) exhibited a combination of promoter-bound CTCF, loss of chromatin accessibility at promoters, and altered interaction with distal regulatory regions (Fig. 5d), indicating that transcriptional repression is frequently associated with multilayered chromatin disruption. Together, these results suggest that *Ctcf* loss preferentially compromises gene expression through direct effects on promoter structure and connectivity. Similarly, most upregulated genes at 7 weeks (80%) exhibited at least one detectable chromatin regulatory alteration (Fig. 5e). These changes were most frequently associated with exclusive direct CTCF binding at promoters (14%), alterations in the number of chromatin interaction partners (10%), and a paradoxical loss of chromatin accessibility at promoters (10%). Notably, a substantial fraction of genes displayed combinatorial regulatory changes, including simultaneous loss of accessibility at promoters and interacting distal regions (10%) or concurrent CTCF promoter binding and altered interaction profiles (7%). This complexity suggests that early transcriptional upregulation following CTCF loss often arises from the combined disruption of local chromatin accessibility and higher-order chromatin architecture. At 13 weeks, the relationship between chromatin changes and transcription was more attenuated. Only 58% of both downregulated (Fig. 5f) and upregulated (Fig. 5g) genes were associated with detectable chromatin regulatory alterations, indicating a partial uncoupling between chromatin state and transcriptional output at later stages or stabilization of changes that occurred at 7 weeks. Among downregulated genes, the dominant event associated with repression of these genes was the CTCF binding at promoters (35%), while changes in interaction with distal regulatory elements represented 11%, either alone or in combination with promoter binding (Fig. 5f). A similar distribution was observed for upregulated genes, with promoter-associated CTCF binding accounting for 30% of cases, followed by changes in interaction with distal regulatory elements alone (11%) and their combination (9%) (Fig. 5g). These data indicate that by 13 weeks more changes in gene expression occur as an indirect consequence of *Ctcf* deficiency and suggest that, over time, compensatory mechanisms may buffer the transcriptional consequences of chromatin accessibility loss, leaving alterations in chromatin topology as a more prominent determinant of gene expression. Given the recurrent association between transcriptional changes and altered chromatin interactions, we next examined how variations in the number of gene promoter-interacting partners correlated with gene expression, by further analyzing pcHi-C and snRNA-seq data. DEGs were classified based on whether they exhibited no change or a decrease/increase (small, medium, or large) in the number of chromatin interactors between control and Ctcf^mKO^ conditions. At 7 weeks, most changes in interaction number were associated with a tendency toward gene downregulation, except for genes showing a large decrease or a small increase in genomic interaction partners (Supp. Fig. 12a). In contrast, at 13 weeks, any change in the number of interacting partners, regardless of direction or magnitude, was associated with a general trend toward gene upregulation (Supp. Fig. 12g). Collectively, these results indicate that *Ctcf* loss exerts temporally distinct effects on gene regulation. Early after depletion, transcriptional changes are tightly linked to direct decrease in chromatin accessibility at promoters and distal regulatory elements, predominantly leading to repression. At later stages, gene expression appears increasingly influenced by global rewiring of chromatin interactions, potentially reflecting adaptive reorganization of the 3D genome that favors transcriptional activation despite persistent *Ctcf* deficiency.

To assess the functional relevance of changes in chromatin interaction profiles, we examined the biological processes enriched among genes stratified by the magnitude of interaction partner changes at 7 and 13 weeks. Genes whose number of genomic interacting partners remained unchanged at 7 weeks were predominantly associated with processes related to synaptic transmission and myotube differentiation and mostly characterized as downregulated genes (Supp. Fig. 12b) that exhibit loss of chromatin accessibility at promoters and non-promoter elements. Genes exhibiting a small decrease in the number of interaction partners were enriched for pathways involved in axon regeneration, including processes such as lamellipodium assembly, as well as responses to leptin signaling (Supp. Fig. 12c). In contrast, genes showing a small increase in chromatin interactions were primarily associated with metabolic processes, particularly those involved in glycogen metabolism and insulin responsiveness, as well as muscle contraction, and mostly referred to downregulated genes (Supp. Fig. 12d). A similar functional signature was observed among genes in the medium increase category (Supp. Fig. 12e), indicating that progressive gains in chromatin connectivity are preferentially associated with downregulation of genes implicated in muscle metabolic and contractile programs. Finally, genes displaying a large increase in the number of interaction partners were significantly enriched for biological processes related to muscle regeneration and function, and immune system regulation, including interleukin-4 production, as well as muscle contraction and cytoskeletal organization and included similar number of down and up-regulated genes (Supp. Fig. 12f). At 13 weeks, DEGs whose number of chromatin interaction partners remained unchanged were enriched for biological processes associated with starvation responses, regulation of apoptotic pathways, activation of adaptive immune responses, and ER stress (Supp. Fig. 12h). This enrichment suggests that, at later stages following *Ctcf* loss, a subset of transcriptional changes occurs regardless of the rewiring of chromatin interactions, and instead reflects sustained cellular stress, metabolic imbalance, and immune activation within myofibers, possibly as indirect consequence of *Ctcf* deficiency. Genes exhibiting a small decrease in the number of interaction partners were predominantly associated with metabolic processes (Supp. Fig. 12i), indicating that even modest reductions in chromatin connectivity can selectively impact metabolic gene networks. In the medium decrease category, enriched biological processes included membrane-associated signaling, ion channel homeostasis, cytoskeletal organization, and metabolic regulation (Supp. Fig. 12j). The convergence of these pathways points to coordinated dysregulation of signaling and structural integrity, processes that are essential for muscle function and adaptability. The simultaneous involvement of ion channel handling and cytoskeletal dynamics further suggests impaired coupling between membrane signaling and intracellular structural responses, possibly reflecting altered response to exposure to physiological perturbations (e.g. contractile or metabolic) of myofiber homeostasis. Genes with a large decrease in interaction partners were enriched for pathways related to lipid biosynthesis and cytoskeletal organization (Supp. Fig. 12k). This pattern indicates that extensive loss of chromatin interactions preferentially disrupts anabolic and structural gene programs, potentially compromising membrane composition, energy storage, and mechanical stability in *Ctcf*-deficient myonuclei. Finally, genes showing a small increase in interaction partners were enriched for biological processes related to complement activation, protein post-translational modifications, and regulation of transcription (Supp. Fig. 12l). These enrichments suggest that gains in chromatin connectivity at 13 weeks may preferentially engage immune-related and regulatory pathways, possibly reflecting compensatory or maladaptive transcriptional responses to chronic chromatin perturbations.

Overall, comparison of biological processes enrichments at 7 and 13 weeks reveals a temporal shift in the functional consequences of chromatin interaction remodeling following *Ctcf* loss. At 7 weeks, changes in chromatin interactions predominantly affected metabolic regulation, muscle contraction, cytoskeletal organization, leading mainly to genes downregulation, indicating an early impact on core muscle functional programs. By 13 weeks, the enriched biological processes increasingly reflected an extension of dysregulation of gene expression to cellular stress responses, immune activation, lipid metabolism, and apoptotic pathways, suggesting a transition from acute functional dysregulation toward chronic stress adaptation and tissue remodeling.

A representative locus of downregulated genes, the *Myh4* locus, is shown in Fig. S13. At 7 weeks, downregulation of *Myh4* in *Ctcf*-deficient myonuclei (Suppl. Figure 13a and b) was associated with loss of chromatin accessibility at *Myh4* promoter (which was bound by CTCF in control myonuclei) and partial reduction of chromatin interactions between *Myh4* promoter and a distal super-enhancer (SE) known to regulate fast-to-slow shift of myosin expression during development(*26*, *63*) (Suppl. Figure 13c). This SE contains multiple peaks of CTCF in control myonuclei coinciding with the chromatin interactions with *Myh4* promoter (Suppl. Figure 13c). By 13 weeks *Myh4* promoter shows continuous absence of chromatin accessibility and complete loss of interaction with the SE, which exhibits a slight increase in chromatin accessibility that did not recover the trend of downregulation of *Myh4* transcription (Suppl. Figure 13c).

While the data described so far indicate that myofiber specific deficiency of *Ctcf* leads to cell autonomous alterations of 3D chromatin structure and gene expression that eventually culminate with myofiber atrophy, snRNA-seq indicates also non-cell autonomous alterations of muscle-resident cells, as an indirect effect of *Ctcf* deficiency (Fig. 4a).

In particular, by week 13, muscles of *Ctcf*^mKO^ mice exhibited a massive infiltration of immune cells that were specifically enriched in pro-inflammatory macrophages, T cells, and dendritic cells (Fig. 6a and b). Heatmap representation of pseudo-bulk differential expression analysis of muscle inflammatory cells shows that most of the changes in gene expression occurred at 13 weeks of age (Fig. 6c). This immune infiltration occurred spontaneously, in the absence of any inflammatory stimulus or trigger, and coincided with the upregulation of various inflammatory genes from type IIB myonuclei of *Ctcf*^mKO^ mice mostly at 13 weeks (shown in Supp. Fig. 7), presumably as a consequence of the combination of chronic activation of ER stress/UPR and 3D chromatin alterations caused by *Ctcf* deficiency(*46–49*). Pseudo-bulk gene ontology of immune cells indicates that while initial changes in the expression of genes regulating B cell proliferation occurred by week 7 (Fig. 6d), more extensive changes in gene expression occurred by week 13 and were related to upregulation of genes implicated in various inflammatory processes, such as expression of cytokines and receptors implicated in the activation of innate and T cell mediated immune response (Fig. 6d). As a result, muscles of *Ctcf*^mKO^ exhibited infiltration by various immune cell types, including macrophages, that were detected in close proximity to myofibers (Fig. 6e-f). This finding suggests that *Ctcf* deficiency in myofibers indirectly activates an immune response, as a non-cell autonomous effect, possibly as consequence of chronic activation of ER stress/UPR combined with myofiber-derived expression of inflammatory genes.

**Fig. 6:**
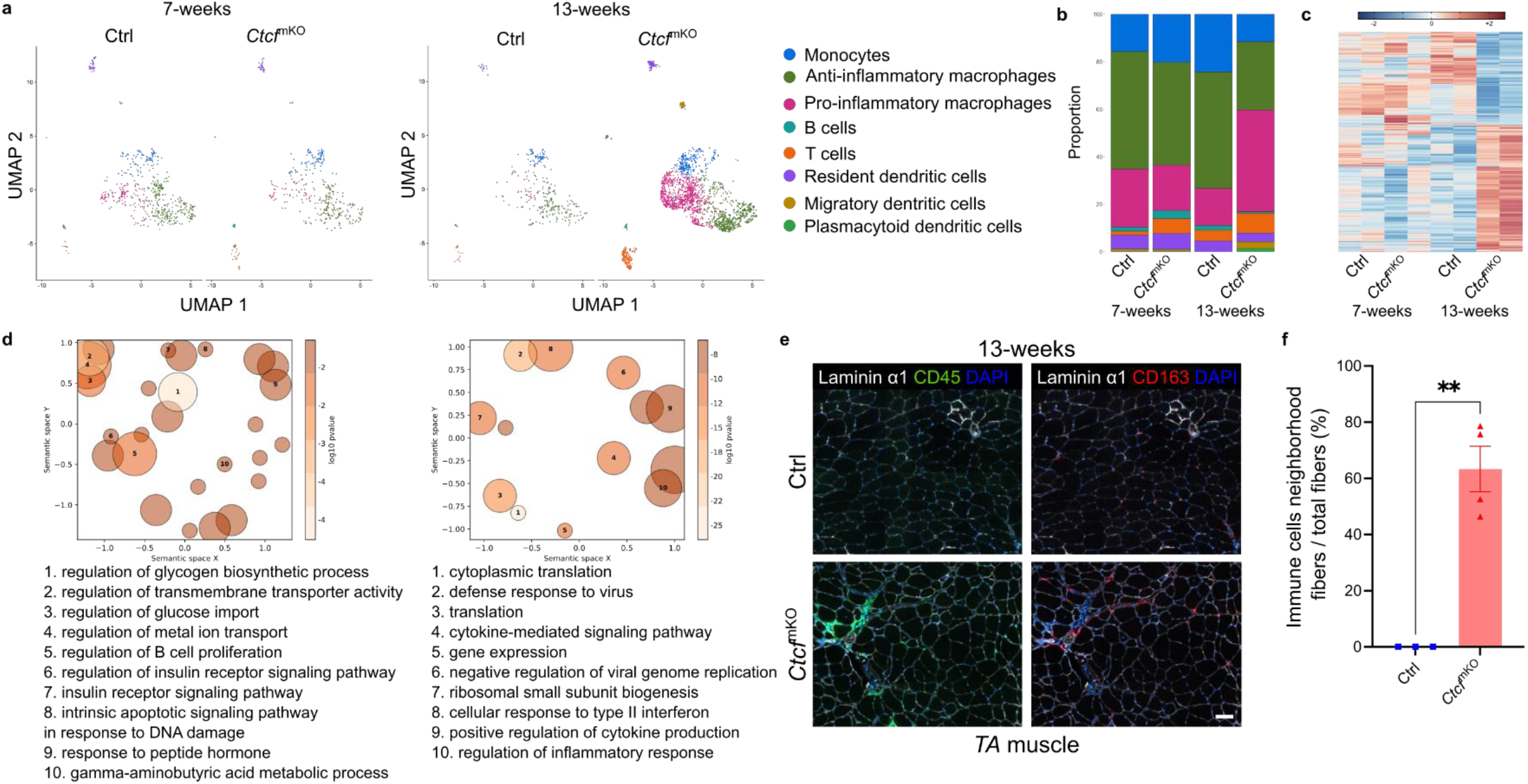
Immune infiltration caused by *Ctcf* depletion in myofibers resembles a myositis-associated atrophic micro-environment. **a**, snRNAseq UMAP of the immune cells metacluster, embedding all conditions: Ctrl and *Ctcf*^mKO^ at 7 and 13 weeks. The dataset is colored and labeled by subcluster. *n* = 2 mice. **b**, Percentage of the immune cells subclusters in Ctrl and *Ctcf*^mKO^ mice at 7 and 13 weeks. Subcluster labeling as in **a**. **c**, Heatmap of differential gene expression generated from the pseudobulk analysis of the immune cells subclusters of Ctrl and *Ctcf*^mKO^ mice at 7 and 13 weeks. *n* = 2 mice. **d**, Top 10 biological processes linked to upregulated genes in immune cells at 7 weeks (left) and 13 weeks (right). Bubbles represent gene ontology terms aggregated by semantic similarity with the GOFigure python package. DEGs defined by log2 FC > 0.25 and *P* value < 0.05. **e**, Representative images of CO-Detection by indexing (CODEX) of transversal section of *TA* muscle of Ctrl and *Ctcf*^mKO^ at 13 weeks, CD45 in green, CD163 in red, Laminin α1 in white and DAPI in blue. Scale bar = 100µm. **f**, Neighborhood analysis of the percentage of muscle fibers surrounded by immune cells in the *TA* muscle of Ctrl (blue squares) and *Ctcf*^mKO^ (red triangles) at 13 weeks. *n* = 3-4 mice; bars are mean±SEM; *P* values of two-tailed unpaired T test.

Overall, our data show that Ctcf deficiency in myofibers induces genome reprogramming toward parallel repression of anabolic genes and activation of stress response (ISR) and inflammation, leading to the spontaneous development of a severe myopathy, with transcriptional features of metabolic and inflammatory myositis.

## Discussion

The results of this study show that disrupting the 3D genome organization in skeletal myofibers, as a consequence of Ctcf deficiency, causes extensive alterations of gene expression that culminate in the spontaneous development of a myopathic syndrome. Thus, this finding reveals in principle the requirement of a continuous maintenance of the integrity of the 3D genome structure in postmitotic, perennial tissues, such as skeletal myofibers.

The results of this study also reveal hitherto unappreciated functional and mechanistic insights into Ctcf-mediated regulation of 3D chromatin structure and gene expression in post-mitotic myofibers. First, they show that *Ctcf* deficiency in post-mitotic myofibers is compatible with developmental myogenesis, but confers susceptibility to spontaneous development of a post-natal myopathic phenotype, indicating a pathogenic “trade-off” whereby genomic alterations caused by *Ctcf* deficiency in post-mitotic myofibers can be tolerated and remain latent during embryonic development at the expense of a spontaneous development of a myopathy upon the exposure to homeostatic perturbations, such as contractile, metabolic or endocrine stimuli encountered in adult life. In this regard, our data reveal an unprecedented causal link between *Ctcf* deficiency and chronic activation of ER stress/UPR and inflammatory genes. Transient activation of ER stress/UPR is typically observed as a physiological consequence of myofiber exposure to post-natal homeostatic perturbations, such as muscle contraction or systemic metabolic and hormonal changes. In contrast, persistent ER stress/UPR is a key event in the pathogenesis of human idiopathic myositis, as it causes the activation of a chronic inflammatory response (*64*, *65*). Interestingly, the phenotype of muscles of *Ctcf*^mKO^ mice by 13 weeks is highly reminiscent of idiopathic inflammatory and metabolic myopathies, as they share salient features, such as muscle atrophy, metabolic and functional impairment, chronic ER stress/UPR, and activation of an inflammatory response (*66–72*). This finding suggests a potential link between CTCF deficiency or haploinsufficiency and the development of human idiopathic myopathies, with chronic activation of ER stress/UPR and inflammation caused by Ctcf-deficiency in myonuclei representing a key pathogenic event. Furthermore, the concomitant downregulation of genes implicated in skeletal muscle development, contraction, metabolism and neurotransmission, and upregulation of genes implicated in myofiber atrophy, proteolysis and catabolism, provides a parallel mechanism that contributes to the myopathic phenotype of *Ctcf*^mKO^ mice.

The extensive alterations in gene expression observed in myonuclei of *Ctcf*^mKO^ mice occur through distinct alterations in chromatin accessibility and interactions between gene promoters and enhancers observed at different stages of post-natal life, as both direct and indirect consequence of Ctcf deficiency, according to the multifunctional role of Ctcf in regulating gene expression(*73–75*). We observed common and fiber-type specific patterns of dysregulated gene expression associated with alterations in chromatin accessibility and promoter-based interactions in Ctcf-deficient myonuclei at distinct stages of myopathy development. At early stages (e.g. 7 week-old *Ctcf*^mKO^ mice) decreased chromatin accessibility at promoters of downregulated genes implicated in myofiber contraction and anabolism, metabolism, adhesion and neuromuscular transmission, appears to occur as a direct consequence of *Ctcf* deficiency and was associated with extensive changes in promoter contacts with distal elements. Conversely, at later stages, upregulation of genes leading to persistent activation of ER stress/UPR and muscle catabolism resulted from global reconfiguration of chromatin structure and connectivity, as an indirect consequence of *Ctcf* deficiency. Interestingly, Ctcf deficiency caused myofiber atrophy despite of the increased levels of the paternally imprinted pro-anabolic gene, Igf2, as previously reported in Thorvaldsen et al.(*76*), and the decreased expression of the pro-atrophic gene, myostatin, indicating that the observed pathogenic phenotype was dominant over the potential compensation from individual genes.

The extensive changes in chromatin structure and gene expression induced by *Ctcf* deficiency in myonuclei lead to a global redistribution and reconfiguration of fiber type, with a progressive loss of type IIB myonuclei, being mostly replaced by myonuclei with gene expression signature predictive of immature, mixed and atrophic fibers. Notably, the residual population of type IIB myonuclei observed in 13 week-old *Ctcf*^mKO^ mice exhibited specific alterations in gene expression suggestive of loss of fiber-type identity and ectopic expression of inflammatory genes. These results reveal the requirement of Ctcf for maintenance of fiber-type identity and transcriptional adaptation of myofibers *in vivo*, through a multilayered control of 3D genome integrity.

Alterations of 3D genome architectures have been associated with the development of human diseases(*16*, *20*, *77*, *78*). Consistently, CTCF deficiency causes developmental abnormalities and has been associated with a wide spectrum of human developmental disorders(*79–81*). Among the CTCF-related human disorders, musculoskeletal defects were annotated in over 50% of the patients analyzed(*27*, *28*). In addition, our previous studies showed that MyoD/CTCF co-bound boundaries of re-wired Insulating Neighborhoods (INs) are hotspots of gene polymorphisms associated with muscular disorders(*29*).

Overall, these data indicate an unprecedented association between *Ctcf* deficiency and susceptibility to the development of myopathies, whereby *Ctcf* dispensability for developmental myogenesis confers vulnerability to develop myopathic syndromes in post-natal life.

Our findings reveal in principle novel biological and mechanistic insights into the role of CTCF as tissue-specific regulator of chromatin structure and gene expression during development. They also encourage future efforts toward establishing whether CTCF deficiency or haploinsufficiency contributes to the pathogenesis of human myopathies as “genetic modifier” of disease severity, thereby extending the spectrum of CTCF-related developmental disorders to human myopathies.

## Limitations of the study

Considering that CTCF-related developmental disorders have been identified based on germline polymorphisms of CTCF, our experimental model of myofiber-specific complete deficiency of *Ctcf* does not completely recapitulate the human conditions of partial systemic CTCF deficiency/polymorphism. However, it suggests that myofiber-specific CTCF deficiency, and related changes in CTCF gene-dosage, might be sufficient to trigger or facilitate the development of myopathies. In this regard, it will be interesting to investigate the potential implication of somatic mutations that might arise in developing myofibers during somitogenesis, as consequences of impaired DNA repair(82, 83) can prompt the generation of myofiber-specific CTCF haploinsufficiency that can instigate the development of myopathic syndromes.

## Acknowledgements

This work was supported by research grants from NIH, R01 GM134712-01 and R01 AR056712 to PLP; French Muscular Dystrophy Association (AFM) postdoctoral grant to JM (AFM 28921, AFM 30566 and AFM 30827); MDA Development Grant to LC (MDA 953791); CIRM postdoctoral training grant to CN; CIRM predoctoral training grant to MN. This study was partly supported by Canadian Institute of Health Research grants to MW, by the Canadian Institutes for Health Research (CIHR) to CC, by the Hong Kong Research Grant Council (GRF16100322, GRF16101524, RFS2425-6S01) to THC, by CAS-Croucher Funding Scheme (CAS24SC02) to THC, by the Innovation and Technology Commission of Hong Kong (ITCPD/17-9) to THC, by the InnoHK initiative of the Innovation and Technology Commission of the Hong Kong Special Administrative Region Government to THC.

This publication includes data generated at the UC San Diego IGM Genomics Center utilizing an Illumina NovaSeq 6000 and Illumina NovaSeq X Plus that were purchased with funding from a National Institutes of Health SIG grant (#S10 OD026929). Flow cytometry was supported by Benji Portillo and Yoav Altman and the SBP Flow Cytometry Shared Resource (RRID:SCR_014854) and snRNA-seq data were generated with the help of Rebecca Porritt and the SBP Genomic core with funding from NCI Cancer Center Support Grant P30CA030199 and Shared Instrumentation Grant S10OD032325. We particularly thank Diana Sandoval, Andy Vasquez, Buddy Charbono and all the animal facility staff members for their help and support on this project.

## Author Contributions

J.M., C.N., L.C. and P.L.P. conceived and designed the experiments. J.M., C.N., L.C., S.H., M.N., J.B. and M.R. performed experiments and data analysis. M.W. provided the Ctcf^fl/fl^ mouse line. S.A. and Y. X. W. provided expertise in CODEX design and analysis. J.M. and P.L.P drafted the manuscript and revised it with contributions from all authors. P.L.P., C.C., M.W., E.M. and T.H.C. conceived the design of the project. P.L.P. supervised and secured funding for the study and wrote the manuscript.

P.L.P. dedicates this work to Brendan, who passed prematurely and will remain forever in the hearth of P.L.P.

## Supplemental information

Figures S1-S13. Table 1.

**Table 1.**
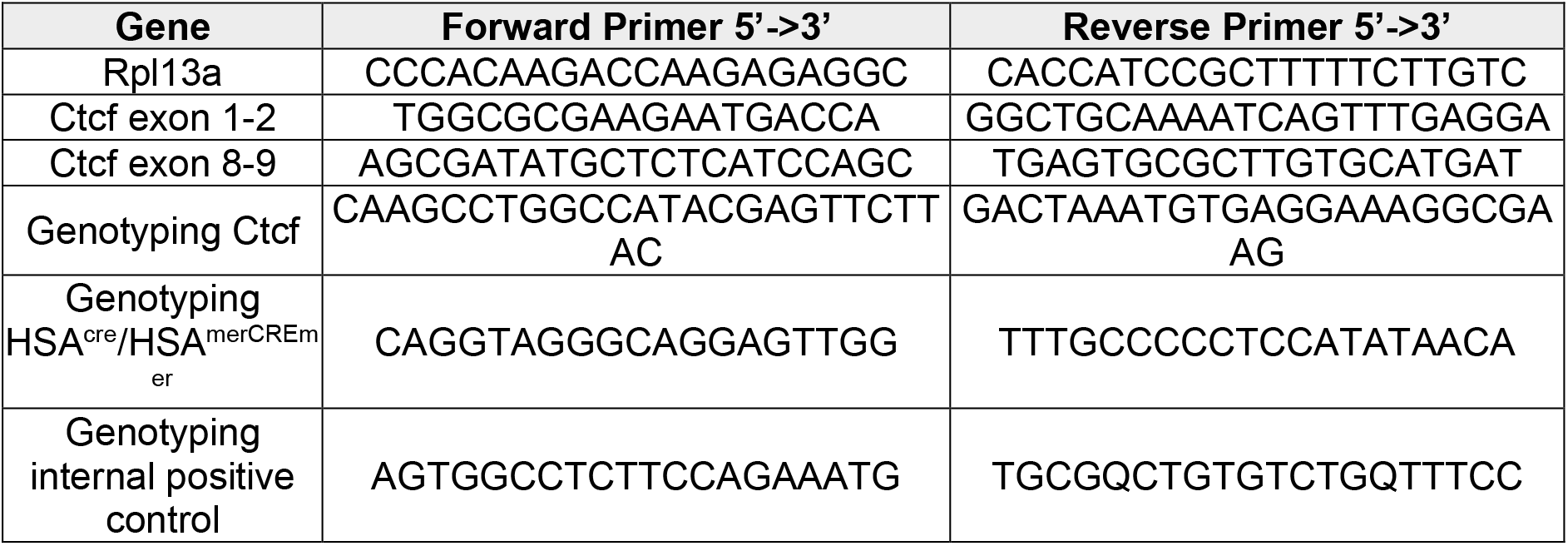
List of primers used.

## Mice

All experiments in this study were performed in accordance with the protocols approved by Sanford Burnham Prebys Medical Discovery Institute Animal Care and Use committee (IACUC) and Canadian animal care organization. B6.Cg-Tg(ACTA1-cre)79Jme/J (HSA^Cre/Cre^) mice were obtained from Jackson Laboratory (Strain #006149). Tg(ACTA1-cre/Esr1*)2Kesr/J (HSA^merCREmer/merCREmer^) was obtained from The Jackson Laboratory (Strain #025750, RRID:IMSR_JAX:025750). *Ctcf^t^*^m1c^ (*Ctcf*^fl/fl^) mice strain was created by *in vitro* fertilization of C57BL/6 N female mice with sperm carrying the *Ctcf*^tm1A(EUCOMM)Wtsi^ mutant allele (Wellcome Trust Sanger Institute, WTSI) at The Centre for Phenogenomics (Toronto, Canada). The Ctcf^tm1A^ allele was flanked by FRT sites, allowing conversion into the conditional *Ctcf^t^*^m1c^ allele containing loxP sites at exon 8 of *Ctcf* gene. HSA^Cre/+^; Ctcf^fl/fl^ (*Ctcf*^mKO^) constitutive muscle specific knockout mice were generated by crossing HSA^Cre/+^; Ctcf^fl/fl^ mice (*Ctcf*^mKO^). HSA^Cre/+^ (Ctrl) and *Ctcf*^mKO^ male mice were used at 7, 9 and 13 weeks-old in this study. HSA^merCREmer/merCREmer^; Ctcf^fl/fl^ (TAM *Ctcf*^mKO^) conditional muscle specific knockout mice were generated by crossing Tg(ACTA1-cre/Esr1*)2Kesr/J and Ctcf^fl/fl^ mice (TAM *Ctcf*^mKO^). Genotypes were routinely confirmed by PCR to verify the experimental cohorts. HSA^merCREmer/merCREmer^ (TAM Ctrl) and TAM *Ctcf*^mKO^ mice were injected with 20mg/kg of body weight of tamoxifen by intraperitoneal injection for 5 consecutive days when they reached the age of 16 weeks. Tamoxifen treated mice were used ar 2 weeks or 4 months after the end of the tamoxifen treatment.

### Tissue preparation

Mice were euthanized and the following tissues were collected. The *Tibialis anterior* (*TA*) and *Extensor digitorum longus* (*EDL*) were isolated separately. *Gastrocnemius, Soleus*, and *Plantaris* muscle were all isolated together and labeled as *GA*. The *TA* muscles were embedded in OCT and quickly frozen by immersion in semi-frozen liquid nitrogen-cooled isopentane and used for immunofluorescence and cross-sectional area analysis. The *EDL* muscles were fixed with cold 0.5% formaldehyde and used for whole muscle immunofluorescence staining. *GA* muscles were used for RNA, protein and nuclei isolation. Whole hindlimb muscles were dissected and used for myonuclei isolation for ChIP. Samplings of the left lobe of the liver were used for protein isolation. Embryos were isolated from euthanized female mice at E14.5 and E16.5 by dating E0.5 as the day after the copulatory plug.

### Hanging test

Mice were subjected to Kondziela’s inverted screen protocol(*84*) to assess muscle endurance. Mice were placed on the center of a wire mesh screen, and a stopwatch was started while the screen was rotated to an inverted position for over 2 seconds. The screen was placed in an inverted position and the duration that the mice held onto the mesh screen until they fell was recorded as a measure of muscular endurance with a set fixed maximum time of 600 seconds.

### *Ex vivo* tetanic force measurement

*TA* muscles from 13 weeks-old mice were isolated and scaled to obtain muscle weight (W_0_ in g) before being placed in Ringer’s buffer at 30°C with 95% O2 and 5% CO2. The tendon was attached to a 300C Dial-Mode Lever System with the Dynamic Muscle Control and Analysis Software following manufacturer recommendations (Aurora Scientific). Optimal muscle length (L_0_ in cm) was measured before gradually adjusting the force to achieve maximal twitch. Peak tetanic force (P_0_ in mN) was measured after successive stimulation of increased frequency from 10 to 200Hz. Specific muscle force (P_1_ mN/cm²) was calculated as followed: P_1_ = P_0_ x (0.44 x L_0_) x 1.06 g/cm^3^ / W_0_.

### Muscle immunostaining

10µm thickness transversal sections were obtained from isolated *TA* muscles in OCT with a Cryostar NX50 cryostat (Thermo Scientific) at -20°C, fixed with formaldehyde 4% in PBS, permeabilized with 0.5% Triton X-100 and blocked with 5% BSA in PBS. Sections were stained with antibody at given dilution (Table 1) O/N at 4°C followed by staining with secondary antibody provided in Table 1 for 1 hour at RT in the dark. Sections were then stained with Hoechst 33258 (Life Technologies) at 2µg/ml for 5 minutes at RT prior to mount in Fluoromount G (Invitrogen, Cat #00-4958-02) Fluorescence microscope was used to acquire images (980 AiryScan, Zeiss) at x10 magnification. Analysis was performed with ImageJ and Biodock softwares.

Fibers’ bundles of prefixed *EDL* muscle were permeabilized with 0.5% Triton X-100 for 1 hour and subsequently incubated in blocking solution (2% BSA, 5% goat serum, 0.5% Triton X-100 in PBS), for 1 hour at RT. Fibers’ bundles were stained with Neurofilament M and SV2A antibodies at given dilution (Table 1), for 48 hours at 4°C. After, samples were incubated twice in blocking solution for 1 hour prior to be incubated with secondary antibodies and conjugated α-bugarotoxin diluted in 4% BSA in 0.5% triton X-100 for 4 hours at RT, protected from light. Samples were subsequently stained with Hoechst 33258 (Life Technologies) at 2µg/ml for 5 minutes at RT prior to several washes in PBS. Samples were then mounted in Fluoromount G (Invitrogen, Cat #00-4958-02) and images were acquired with confocal microscope (980 AiryScan, Zeiss), ×20 magnification and 0.5µm z-stack interval. Images are presented as maximum orthogonal intensity projections of the z-stacks, using Image J software.

### CODEX multiplexed immunofluorescence imaging analysis

Multiplex imaging (CODEX, co-detection by indexing) was performed on 10µm *TA* transversal sections according to the manufacturer’s instructions (Akoya Biosciences) with modified staining steps. Briefly, tissue sections were mounted on 0.2% gelatin coated coverslips. Tissue sections were thawed in hydration buffer, fixed with 1.6% formaldehyde for 10 minutes and stained with the mix containing the entire panel of custom-conjugated lectins and antibodies overnight at 4°C. After washing, coverslips were fixed again with 1.6% formaldehyde for 10 minutes, followed by a final fixation step with 200mg/ml BS3 (bis(sulfosuccinimidyl)suberate; Thermo Fisher) in anhydrous DMSO (Sigma) for 20 minutes. The coverslips were again washed and stored at 4°C until CODEX imaging. Staining mix included the CODEX antibody panel defined in Table 1, in a buffer solution together with 5µL of each of 1mg/ml mouse IgG (Sigma), 1mg/ml rat IgG (Sigma), Sheared salmon sperm DNA (10mg/ml in H_2_O; Thermo Fisher), Mixture of non-modified CODEX oligonucleotides at a final concentration of 0.5mM in TE buffer (IDT) solutions. CODEX was performed on a Phenocycler Open system (Akoya) on a Keyence BZ-810 automated microscope. Fluorescent probes (5µL of a 10mM stock in 250µL of plate buffer) for the matching antibody barcode were loaded into a 96-well plate as per Phenocycler protocol. For each CODEX experiment, Ctrl sample and *Ctcf*^mKO^ samples were sectioned on the same coverslip, stained and imaged together.

Imaging data was processed using CRISP-CODEX processor(*85*) to register (across image tiles and imaging cycles), deconvolve, and generate stitched extended-depth-of-field images of deconvolved optical slices of the muscle sections after background subtraction against autofluorescence.

### Western blot

Samples of *GA* muscles and liver were lysed in Lysis Buffer (20mM Tris-HCl pH 7.5, 0.15M NaCl, 1% Triton X-100, 5mM EDTA pH 8.0, 0.1M NaF, 2mM Na-Pyrophosphate, 1mM PMSF and 1X protease and phosphatase inhibitors). Protease inhibitor was purchased from Roche (COmplete mini, Cat #11836153001) and Pierce Phosphatase inhibitor (Thermo Scientific, Cat #A32957). Protein concentration was measured using BCA protein assay kit (Thermo Scientific, Cat #23227). For each sample, 30μg of proteins were diluted in loading solution and run on 10% tris-glycine gel (Invitrogen, Cat #XP00100B0X) and transferred on PVDF membranes (Invitrogen, Cat #IB24002). As loading control, membranes were stained for 20 minutes at RT with Ponceau S (0.2% Ponceau S, 3% Acetic Acid in H_2_O). After three washes in TBST (20 mM Tris-HCl pH7.6, 0.15M NaCl with 0.1% Tween 20), the membrane was blocked with 5% skim milk (RPI, Cat#M17200) in TBST for 2 hours at RT. The membrane was incubated with the listed primary antibodies (Table 2) in 5% skim milk in TBST O/N at 4°C. The membrane was washed in TBST three times before to be incubated 2 hours with anti-rabbit IgG HRP (Cell Signaling, Cat #7074S) or anti-mouse IgG HRP (Cell Signaling, Cat #7076S). The membrane was washed two times with TBST and once with TBS before detection with SuperSignal^TM^ West Femto Maximum Sensitivity Substrate (Thermo Scientific, Cat #34095).

### mRNA expression analysis

Total RNA was extracted using Quick-RNA Microprep kit (Zymo Research, Cat#R1051) following manufacturer’s recommendation. RNA concentration was measured on Qubit (Invitrogen). 100-500 ng of RNA was reverse transcribed using High-Capacity cDNA Reverse Transcription Kit (Applied Biosystems, Cat#4368813). Real-time quantitative PCR (qPCR) was performed using Power SYBR Green Master Mix (Life Technologies) following manufacturer’s indications. Expression was normalized to *Rpl13A* using 2^−DDCt^ method.

### Nuclei processing for snRNA-seq

For each replicate, 1 *GA* muscle of 7 or 13 week-old mice was dissected and flash frozen in liquid nitrogen, and stored at -80°C. Muscles were thaw and minced in cold lysis buffer (10mM Tris-HCl, 10mM NaCl, 3mM MgCl_2_, 0.1% IGEPAL-CA630, 0.1% Tween-20, 1% BSA, 0.15U/µl RNAse inhibitor Protector (Sigma, Cat#3335402001), 0.15U/µl RNAseOUT (Thermo Fisher, Cat#10777019), 0.15U/µl Superase RNAse inhibitor (Invitrogen, Cat#AM2696), 0.15U/µl Watchmaker RNAse inhibitor (Watchmaker Genomics, Cat#7K0088-500UL) in DEPC water). After mincing, 6ml of cold lysis buffer was added to the tissue and incubated in ice for 10 minutes. The lysate was dounced with 8 strokes of a loose pestle in a 8ml dounce homogenizer and then diluted with 8ml of wash buffer (10mM Tris-HCl, 10mM NaCl, 3mM MgCl_2_, 2% BSA, 0.15U/µl RNAse inhibitor Protector, 0.15U/µl RNAseOUT, 0.15U/µl Superase RNAse inhibitor, 0.15U/µl Watchmaker RNAse inhibitor in DEPC water). The diluted homogenate was subsequently filtered with 100, 70 and 40µm cell strainers. Nuclei were pelleted at 500G for 5 minutes at 4°C at slow acceleration and deceleration settings before to be resuspend in 300µl of cold resuspension buffer (1% BSA, 1X PBS, 0.2U/µl RNAse inhibitor Protector) and stained with 7AAD (Sigma, Cat#SML1633) for 15 minutes. Following staining, 700µl of resuspension buffer was added before to spin down the nuclei with similar settings. A last wash was performed with 500µl of resuspension buffer before to resuspend the nuclei in 300µl of resuspension buffer for FACS sorting. Nuclei were sorted for 7AAD^pos^ with a BD FACSAriaII in 500µl of collection buffer (2% BSA, 1X PBS and 0.4U/µl RNAse inhibitor Protector), spun down at 350G and 4°Cand resuspended in 50µl of resuspension buffer. Nuclei were then counted and diluted to obtain a recommended concentration of 12 000 nuclei/sample for optimal usage of the 10X Genomics 3’ v3 kit. Libraries were constructed following the manufactured recommendation and sequenced with Element Biosciences AVITI Sequencer 2×75bp at 30 000 reads/nuclei.

### Myonuclei isolation for myonuclei bulk ATAC and Promoter Capture HiC

For each replicate, nuclei were isolated from 2 *GA* muscles following the previously described protocol. After filtration in cell strainers, nuclei were enriched for myonuclei using the specific myonuclear marker PCM1. Nuclear suspension was incubated with 0.22µg/ml of rabbit anti-PCM1 antibody (Sigma, Cat#HPA023370-100ul) diluted in staining buffer (5% BSA, 0.2% IGEPAL-CA630, 1mM EDTA in 1X PBS) for 1 hour at 4°C on a rotator. Nuclei were spun down at 500G for 5 minutes at 4°C and washed twice with 1ml of MACS buffer (2% BSA, 0.5mM EDTA in 1X PBS) before being resuspended in beads buffer, composed of 160µl of MACS buffer and 40µl of secondary anti-rabbit IgG microbeads (Miltenyi Biotec, Cat#130-048-602), and incubated for 15 minutes at 4°C. Nuclei were next washed twice with MACS buffer and myonuclei were enriched using Miltenyi recommendation for positive selection of LS column (Miltenyi Biotec, Cat#130-042-401).

### RNA-seq library preparation

Equal inputs of total RNA (10-100 ng) were used to generate stranded total RNA libraries for sequencing using the Illumina® Stranded Total RNA Prep, Ligation with Ribo-Zero Plus (Illumina, Cat #20040525). ERCC RNA Spike-in mix was added at the beginning of the protocol according to manufacturer’s instruction (Ambion, Cat#4456740). Libraries were sequenced at a depth of ∼50 million reads per library on NOVASeqS4 PE 2×100 at the IGM UCSD.

### ATAC-seq library preparation

100 000 freshly collected myonuclei were used to perform an ATAC-seq library preparation using the ATAC-seq Kit (Active Motif, Cat #53150) according to manufacturer’s protocol. Libraries were sequenced at a depth of ∼200 milions reads per library on NOVASeqS4 PE 2×100 at the IGM UCSD.

### Promoter Capture HiC and library preparation

Promoter Capture HiC experiments were performed using the Arima HiC+ kit (Arima Genomics, Cat#A101020), Accel-NGS 2S Plus DNA Library Kit (Swift Biosciences, Cat #21096) and Swift Indexing Kit (Swift Biosciences Cat #26148, 26248) according to manufacturer’s protocol. Briefly, 1.5 million myonuclei were resuspended in 5ml of MACS buffer and fixed with 2% formaldehyde for 10 minutes at RT followed by quenching with STOP solution1 according to Arima HiC+ protocol. Digestion was performed O/N at 37° C. 1 µg of purified HiC libraries were hybridized at 65° C to Arima Mouse Promoter Panel (Arima genomics, Cat#A311026) and enriched with Streptavidin beads according to Arima Capture-HiC+ protocol. Obtained libraries were sequenced on NOVASeqS4 PE 2×100 at the IGM UCSD at a depth of ∼300 million reads per library.

### CTCF ChIP-seq library preparation

For each replicate, nuclei were isolated from 4 *GA* muscles of 8 to 12-weeks old wild type C57BL/6 following the previously described protocol. After filtration in cell strainers, nuclei were fixed with 1% formaldehyde for 5 minutes at RT. They were next quenched with 125mM glycine for 5 minutes at RT and subsequent 10 minutes in ice. After one wash with 1X PBS, the nuclei were enriched for myonuclei using PCM1 antibody as previously described. For each ChIP-seq replicate, 6×10^6^ myonuclei were resuspended in lysis buffer (50mM Tris-HCl pH 8.0, 150mM NaCl, 5mM EDTA pH 8.0, 0.5% SDS, 0.5% NP-40, 1mM PMSF and 1x protease inhibitor). Chromatin was sheared using Bioruptor Pico sonication device (Diagenode), for 30s sonication and 30s of rest for 25 cycles, to obtain an average DNA fragment length of 200-500bp. 15µg of sonicated chromatin were diluted in 500µl RIPA Dilution Buffer (50mM Tris-HCl pH 8.0, 150mM NaCl, 1mM EDTA pH 8.0, 0.1% Sodium Deoxycholate, 0.7% NP-40, 1mM PMSF and 1x protease inhibitor), and precleared for 3 hours at 4° C with Protein A/G dynabeads. In parallel, 10µg of CTCF antibody (Cell Signaling, Cat #3418S), or 1µg of rabbit IgG (Santa Cruz, Cat #sc-2027) were prebound to Protein A/G dynabeads in 500µl of PBS containing 5 µg/ml BSA, for 3 hours at 4° C. Beads bound antibody were mixed with precleared chromatin and incubated O/N at 4° C on a rotator. Chromatin bound fraction was washed with RIPA washing buffer (50mM Tris-HCl pH 8.0, 150mM NaCl, 1mM EDTA pH 8.0, 0.5% Sodium Deoxycholate, 1% NP-40) for 4 times, followed by LiCl washing buffer (10mM Tris-HCl pH 8.0, 250mM LiCl, 1mM EDTA pH 8.0, 1% Sodium Deoxycholate, 1% NP-40), and TE buffer (10mM Tris-HCl pH 8.0, 1mM EDTA), each wash for 10 minutes at 4° C. Chromatin was then eluted in elution buffer (1% SDS in TE buffer) at 65° C for 6 hours 600 RPM rotation. Precipitated material and 10ng of input were used as input for library preparation using NEBNext® Ultra™ II DNA Library Prep Kit (NEB, Cat #E7645L) and NEBNext® Multiplex Oligos (NEB, Cat #E7600S) according to manufacturer’s protocol. Obtained libraries were sequenced on Element Biosciences AVITI Sequencer 2×75bp at a depth of ∼100 million reads per library.

### snRNA-seq data preprocessing, dimensionality reduction, and visualization

Data were aligned to the mm10-3.0.0/GRCm38 version of the mouse genome using CellRanger (v8.0.1; 10x Genomics; https://github.com/10XGenomics/cellranger/releases) with parameter --include-introns true. Ambient RNA transcripts were removed with Cellbender (v0.3.0; https://github.com/broadinstitute/CellBender)(86) and the doublet score was calculated with Scrublet (v0.2.3; https://github.com/swolock/scrublet)(87) at a prediction rate of 8%(*87*). Samples were analyzed with Seurat (v5.2.1; https://github.com/satijalab/seurat/releases) package in R (v4.3.2; https://cran.r-project.org/bin/windows/base/old/)(*88*). The samples contained an average of 10,480 nuclei and 1,140 genes. They were merged and filtered for quality control to remove nuclei with less than 200 genes, more than 2,500 genes, a Scrublet-derived doublet score higher than 0.2 and a percentage of mitochondrial DNA higher than 5%. Dimensional reduction was performed using Principal Component Analysis (PCA) on 50 vectors of similar expression. Next, a first batch correction was performed using Seurat’s SCTransform function with parameter vars.to.regress using the counts of mitochondrial genes, the number of genes, the cell cycle phases and the Scrublet-derived doublet score. A second batch correction was then performed for the 3 batches of nuclei preparation using Seurat’s function IntegrateLayers with HarmonyIntegration method, SCT as normalization method and PCA as the original reduction. FindNeighbors, FindClusters and RunUMAP were based on the optimal number, 35 PCs over 50 PCA, choose with the elbow plot function. FindClusters was run with parameters, Resolution = 0.2, Algorithm = 3. RunUMAP was performed using HarmonyIntegration reduction. PrepSCTFindMarkers and FindAllMarkers were used to find the 10 most differentially expressed genes in each cluster to define their identity. Each metacluster was then subset and reanalyzed with FindNeighbors, FindClusters and RunUMAP to defined subclusters. Through FindAllMarkers, an identity to the subcluster was given, and subset function was used to remove contaminants clusters. Finally, all layers and regression were removed from each cluster before using Seurat function merge, to merge all subclusters into a final object. Layers corresponding to the 3 batches of nuclei preparation were added to the final object and it was processed with all previously presented function and parameters for further analysis. To characterize myonuclei trajectories, pseudo-time analysis was performed using the “Monocle3” R package (version 1.4.26; https://cole-trapnell-lab.github.io/monocle3/)(*89–92*). The earliest principal node has been set to the canonical fiber type for each of the analyses.

### Myonuclei pseudo bulk snRNA-seq data analysis

Nuclei belonging to myofibers were subset from the complete dataset Seurat object with the subset function, RNA counts layers collapsed through the JoinLayers function, and the object was split again by time point (7 and 13 weeks). Counts were then aggregated across sample groups with the aggregate.Matrix function (with fun = “sum”). Samples with zero cell contributing to the cluster were removed, and the final data frame (capturing metadata specific to cluster) was joined to the metadata object through the plyr package function join. A DESeq object was made using the DESeq2 (v1.34.0; https://bioconductor.org/packages/release/bioc/html/DESeq2.html)(93) function DESeqDataSetFromMatrix, principal component analysis was carried out with the plotPCA function, and differential analysis was performed with the DESeq function, with differentially expressed genes considered with a threshold of p-val < 0.05 and |log2 fold change| > 0.25.

### RNA-seq data analysis

Data were checked for quality with FASTQC (v0.11.9; http://www.bioinformatics.babraham.ac.uk/projects/fastqc/), reads were trimmed with Trimmomatic (v0.39; http://www.usadellab.org/cms/?page=trimmomatic)(94) to eliminate low quality bases and adapters (parameters: PE -phred33 ILLUMINACLIP:NexteraPE-PE.fa:2:30:10:2:true MAXINFO:40:0.1 MINLEN:45), and aligned to the mm10 Ensembl (November 2019) version of the mouse genome, with STAR (v 2.7.3a; https://github.com/alexdobin/STAR)(95), with parameters: --runMode alignReads -- runThreadN 12 --readFilesCommand zcat --outSAMtype BAM SortedByCoordinate -- quantMode GeneCounts. Counts data (from STAR output in column 2, based on library preparation strandedness: unstranded) from all conditions were filtered based on their raw count, keeping genes where the sum of the counts for all samples was higher than 10. Size factors were calculated using the R environment (v4.1.3)(*96*) package DESeq2 estimateSizeFactors function, then normalized and logged with the rlog function. DESeq2 was also used to perform Principal Component Analysis (PCA) and differential gene expression analysis (significance threshold: BH FDR<0.05). Differential expression analysis results were visualized with the heatplot function (parameters: method=“complete”, labRow= FALSE, dend=“row”, returnSampleTree=F, zlim=c(-5,5)) from the made4 (v1.68.0)(*97*) from Bioconductor package (https://datacarpentry.github.io/genomics-r-intro/04-bioconductor-vcfr.html)(*98*), after expression values from the DESeq2 object were turned into a matrix of Z-scores. Gene Ontology was performed on down- and up-regulated genes separately, using the gene symbols as input in EnrichR (https://maayanlab.cloud/Enrichr/)(99), and gene sets repositories GO Biological Processes 2023 and Descartes Cell Types and Tissue 2021. Top10 gene ontology terms shown in figure panels were ordered by pvalue. Gene Ontology bubble plots to summarize top50 gene ontology terms, were generated with the Python (v.3.8.1; https://www.python.org/downloads/release/python-381/)(100) GO-Figure (https://gitlab.com/evogenlab/GO-Figure)(*101*) script (parameters: -j standard -c log10-pval - e 100 -si 0.5 -s members -g single -n bpo --font_size small -q svg).

### ATAC-seq data analysis

Data were checked for quality with FASTQC, reads were trimmed with Trimmomatic to eliminate low quality bases and adapters (parameters: PE -phred33 ILLUMINACLIP:NexteraPE-PE.fa:2:30:10:2:true MAXINFO:40:0.1 MINLEN:45), aligned to the mm10 Ensembl (November 2019) version of the mouse genome with Bowtie2 (v2.3.5.1)(*102*) (parameters: --no-unal --local --very-sensitive-local --no-discordant --no-mixed --dovetail --phred33), sorted and converted into BAM format with the SAMtools sort function. Peak calling was performed with the Macs2 (v2.2.6; https://pypi.org/project/MACS2/)(103) callpeak function (parameters: -f BAMPE -g hs -B -q 0.05 --nomodel --shift -100 --extsize 200), filtered for black-list regions with the BEDtools suite (v2.29.2; https://bedtools.readthedocs.io/en/latest/)(104) intersect (parameters: -v), and sorted with the sort function (parameters: -k8,8nr). Irreproducible Discovery Rate (IDR) was calculated with the idr function (v2.0.3; https://github.com/nboley/idr)(105) (parameters: --input-file-type narrowPeak --plot --only-merge-peaks) to retain a single peaks list from each condition. Differential chromatin accessibility analysis was carried out in R, with the DESeq2 package. The R package ChIPQC (v1.30.0; https://bioconductor.org/packages/release/bioc/html/ChIPQC.html)(106), (GetGRanges function was used to import peak lists from each biological replicate, a consensus peak list was calculated for all conditions using the IRanges (v2.28.0; https://bioconductor.org/packages/release/bioc/html/IRanges.html)(107),function reduce, keeping peaks that were called in at least two samples. Counts for each peak in each biological replicate were quantified with the Rsubread (v2.8.2; https://bioconductor.org/packages/release/bioc/html/Rsubread.html)(108) featureCounts function (parameters: isPairedEnd= T, countMultiMappingReads= F, maxFragLength= 100). Counts were normalized and logged with the DESeq2 function rlog; DESeq2 was also used to perform Principal Component Analysis (PCA), and differential chromatin accessibility analysis (significance threshold: BH FDR<0.05). Differentially Accessible Regions (DARs) at gene promoter regions were retrieved with the IRanges function promoters (parameters: TxDb.Mmusculus.UCSC.mm10.knownGene, 1000, 200). Promoter DARs were visualized with the made4 function heatplot (parameters: method=“complete”, labRow= FALSE, dend=“row”, returnSampleTree=F, zlim=c(-5,5)), after expression values from the DESeq2 object were turned into a matrix of Z-scores. Due to R visualization limits, only promoters with |log2FC|>1.5 were plotted. Motif analysis was carried out with the HOMER (v4.11; http://homer.ucsd.edu/homer/index.html)(109), findMotifsGenome function (parameters: mm10 -size given -nomotif).

### ChIP-seq data analysis

CTCF ChIP-seq data were checked for quality with FASTQC and aligned to the mm10 Ensembl (November 2019) version of the mouse genome with Bowtie2 (parameters: --very-sensitive-local). Peak calling was performed with the Macs2 callpeak function (parameters: - g mm -B --nomodel --SPMR -p 0.05), using input chromatin as control.

### Promoter capture Hi-C data analysis

pcHiC data were analyzed using the Arima Genomics proprietary script Arima-CHiC-v1.5.sh, which comprises the following analysis steps: reads truncation (based on the restriction enzymes used), alignment, filtering and PCR removal through the HiCUP pipeline(https://stevenwingett.github.io/HiCUP/)(110), the function bam2chicago, and ChICAGO(https://github.com/dovetail-genomics/chicago)(111) for loop calling, at a resolution of 5kb. The script was launched with the following parameters: -A bowtie2-2.3.5.1/bowtie2 -X ARIMA_Capture_HiC_Settings/mm10/reference/mm10 -d ARIMA_Capture_HiC_Settings/mm10/reference/Digest_mm10_Arima.txt -H HiCUP-0.8.0 -C chicagoTools -b ARIMA_Capture_HiC_Settings/mm10/mouse_GW_PC_S3207063_S3207103_mm10.uniq.b ed -R ARIMA_Capture_HiC_Settings/mm10/5kb/mm10_chicago_input_5kb.rmap -B ARIMA_Capture_HiC_Settings/mm10/5kb/mm10_chicago_input_5kb.baitmap -D ARIMA_Capture_HiC_Settings/mm10/5kb/ -O mm10 -r 5kb. A merged peak list for each condition was obtained with the ChiCAGO auxiliary function makePeakMatrix.R, with parameter –twopass, and together with the counts from ChICAGO (stored in the chicinput files), and the data stored in the ChICAGO RDS objects, differential interactions were identified with Chicdiff(*112*), with parameters settings= list(“saveAuxData“=T, “score“=3).

### Genome-wide tracks generation and visualization

Genome-wide tracks of RNA-seq signal in BEDGRAPH format (forward and reverse) were generated with the DeepTools (v3.5.4; https://github.com/deeptools/deeptools/releases)(113), function bamCoverage (parameters: --normalizeUsing CPM --effectiveGenomeSize 2652783500 --binSize 100 --filterRNAstrand forward or reverse --outFileFormat bedgraph -- scaleFactor $sizeFactor) using as scale factors the size factors calculated with DESeq2 (where spike-in reads were used as control genes), filtered for non-standard chromosomes, then joined with the Unix function cat, sorted with the sort function (parameters: -k1,1 -k2,2n), compressed with bgzip and indexed with tabix (parameters: -p bed) to be visualized within the Washington University (WashU) Epigenome Browser (http://epigenomegateway.wustl.edu/browser/)(114). Tracks in BIGWIG format for ATAC-seq, CUT&RUN, and ChIP-seq data were generated with the DeepTools function bamCoverage (parameters: --normalizeUsing CPM --effectiveGenomeSize 2652783500 --binSize 10 -- extendReads 300 --ignoreDuplicates). The DeepTools function bigwigAverage was used to generate an average signal profile of ATAC-seq for each condition, while also filtering out signal from blacklist regions. Tracks were visualized either within the WashU epigenome browser or in tornado and aggregate signal plots generated with the DeepTools functions computeMatrix scale-regions (parameters: --startLabel Start --endLabel End -- beforeRegionStartLength 3000 --afterRegionStartLength 3000 --skipZeros -- missingDataAsZero); computeMatrix reference-point (parameters: --referencePoint center -- beforeRegionStartLength 3000 --afterRegionStartLength 3000 --skipZeros -- missingDataAsZero); plotHeatmap (parameters: --xAxisLabel “” --yAxisLabel “Coverage” -- heatmapHeight 12 --yMin 0 --refPointLabel “peak center”) with --sortUsingSamples 1; plotProfile (parameters: --yAxisLabel “Coverage” --yMin 0 --refPointLabel “peak center” -- perGroup --startLabel Start --endLabel End).

### Data integration

Overlaps of different genomic intervals was performed with the BEDtools function intersect, to retain a unique list of overlapping features (parameters: -wa -u), or to integrate different features to be processed within R for quantification (parameters: -wao), or to select only non-overlapping features (parameters: -v). Promoter regions were defined as -1000/+200 bp from the gene TSS, according to GENCODE.vM10.annotation.gff3 and RefSeq mm10 (from UCSC Table Browser), restricting the analysis to protein coding and lincRNA (GENCODE) and known genes (RefSeq). Partition of ATAC peaks into genomic features (intergenic or intragenic: upstream, -UTR, Exon, Intron, -UTR, Downstream) was carried out according to PAVIS (available online at https://manticore.niehs.nih.gov/pavis2/)(115).

---

**Fig. S1.**
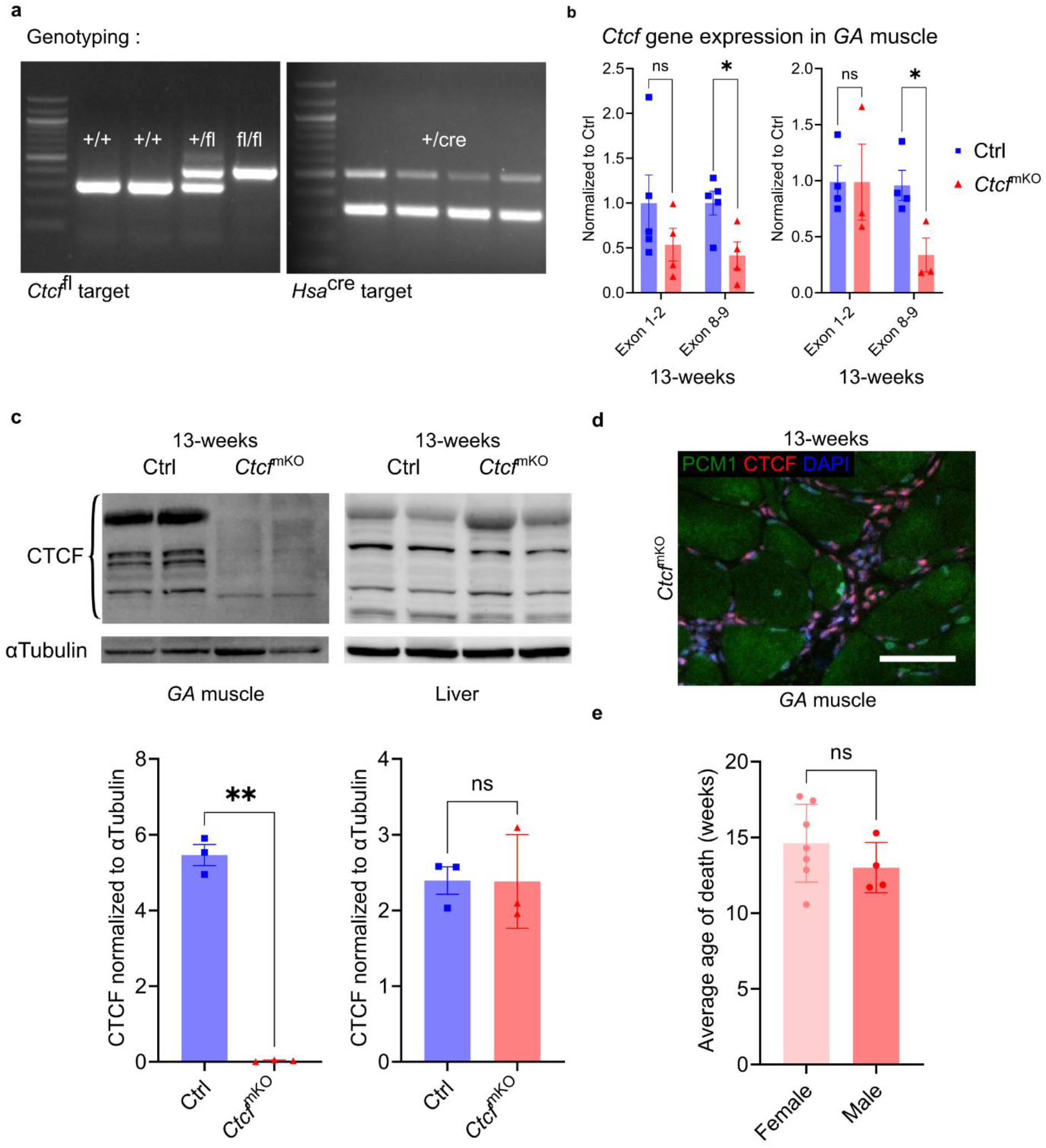
CtcfmKO mice exhibit loss of Ctcf gene and protein expression in muscle tissue. a, Representative images of Ctcffl and Hsacre genotyping. b, RT-qPCR of Ctcf gene expression with primers against exon 1-2 and exon 8-9 in whole GA muscle of Ctrl (blue squares) and CtcfmKO (red triangles) mice of 7 and 13 weeks. n = 3-5 mice; bars are mean±SEM; P values from two-tailed unpaired T test. c, Western blot of CTCF protein in GA muscle (left) and liver (right) of Ctrl (blue squares) and CtcfmKO (red triangles) mice at 13 weeks, western blot membrane (upper panel) quantification to αTubulin (lower panel). n = 3; bars are mean±SEM; P values from two-tailed unpaired T test. d, Representative image of the transversal section of TA muscle of CtcfmKO at 13 weeks, PCM1 myonuclei marker in green, CTCF in red and nuclei in blue. Scale bar = 50µm. e, Average life expectancy of CtcfmKO female (pink) and male (red) mice in weeks. n = 4-7; bars are mean±SEM; P values from two-tailed unpaired T test. *P<0.05; *** P<0.005; ns = non significant.

**Fig. S2.**
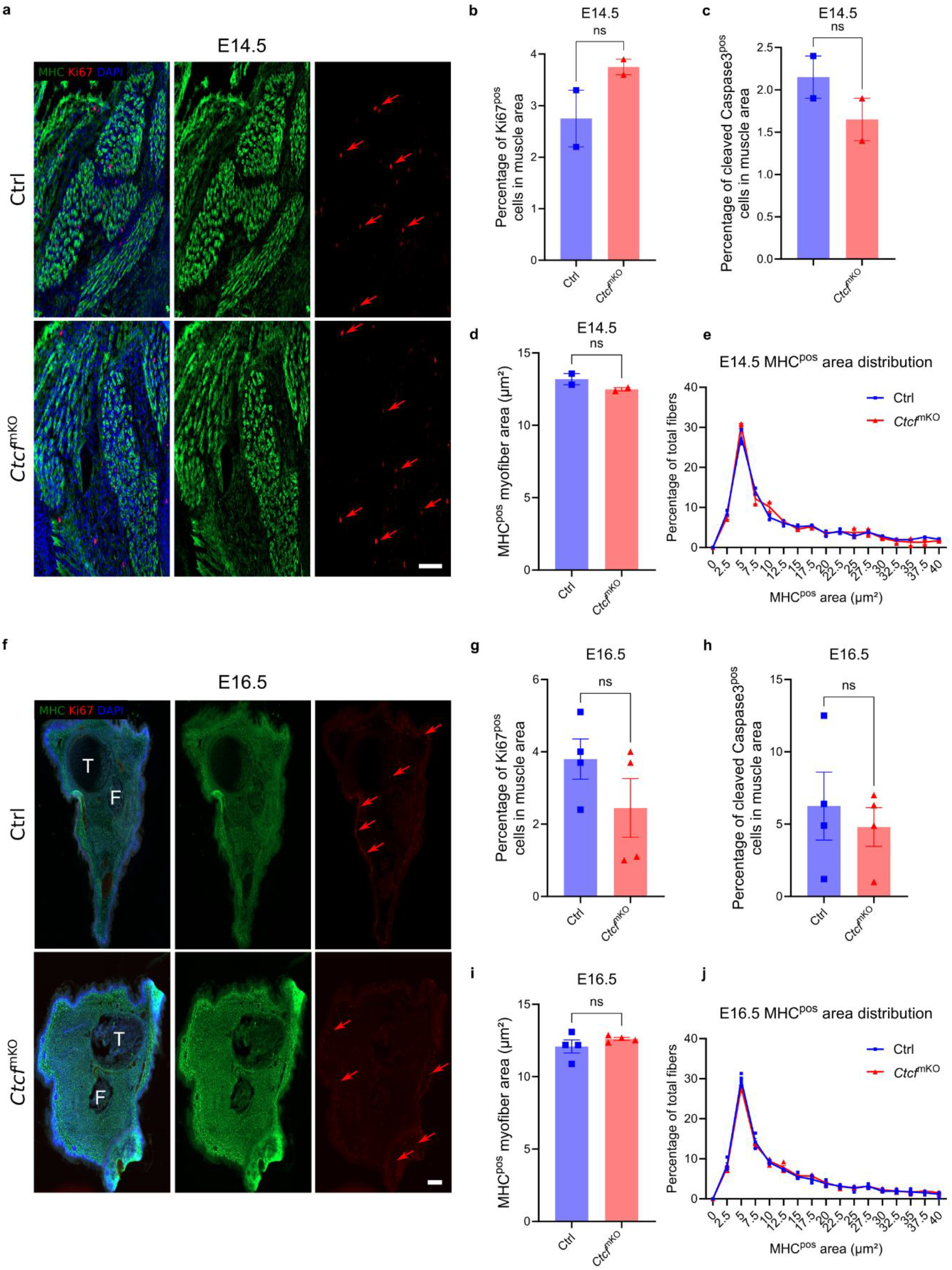
The adult phenotype does not result from late embryogenic defects. a, Representative images of immunostaining of transversal section of body wall of Ctrl and CtcfmKO embryo at E14.5, all myosin heavy chains (MHC) in green, Ki67 in red and DAPI in blue. Red arrows indicate Ki67 positive cells. Scale bar = 100µm. b, Percentage of Ki67 positive cells in the E14.5 embryonic muscles from a. Ctrl (blue) and CtcfmKO (red). n = 2 embryos; bars are mean±SEM; P values from two-tailed unpaired T test. c, Percentage of cleaved caspase 3 positive cells in the embryonic muscles from a. n = 2 embryos; bars are mean±SEM; P values from two-tailed unpaired T test. d, Quantification of the percentage of area of MHC-positive myofibers in Ctrl and CtcfmKO from a. n = 2 embryos; bars are mean±SEM; P values from two-tailed unpaired T test. e, Frequency of the MHC-positive myofiber per area distribution. n = 2 embryos. f, Representative images of immunostaining of transversal section of hindlimb of Ctrl and CtcfmKO embryo at E16.5, MHC in green, Ki67 in red and DAPI in blue. Red arrows indicate Ki67 positive cells. T = Tibia; F = Femur; scale bar = 200µm. g, Percentage of Ki67 positive cells in the embryonic muscles from f. n = 3 embryos; bars are mean±SEM; P values from two-tailed unpaired T test. h, Percentage of cleaved caspase 3 positive cells in the embryonic muscles from f. n = 3 embryos; bars are mean±SEM; P values from two-tailed unpaired T test. i, Quantification of the percentage of area of MHC-positive myofibers in Ctrl and CtcfmKO from f. n = 3 embryos; bars are mean±SEM; P values from two-tailed unpaired T test. j, Frequency of the MHC-positive myofiber per area distribution from f. n = 3 embryos; bars are mean±SEM; P values from two-tailed unpaired T test. ns = non significant.

**Fig. S3.**
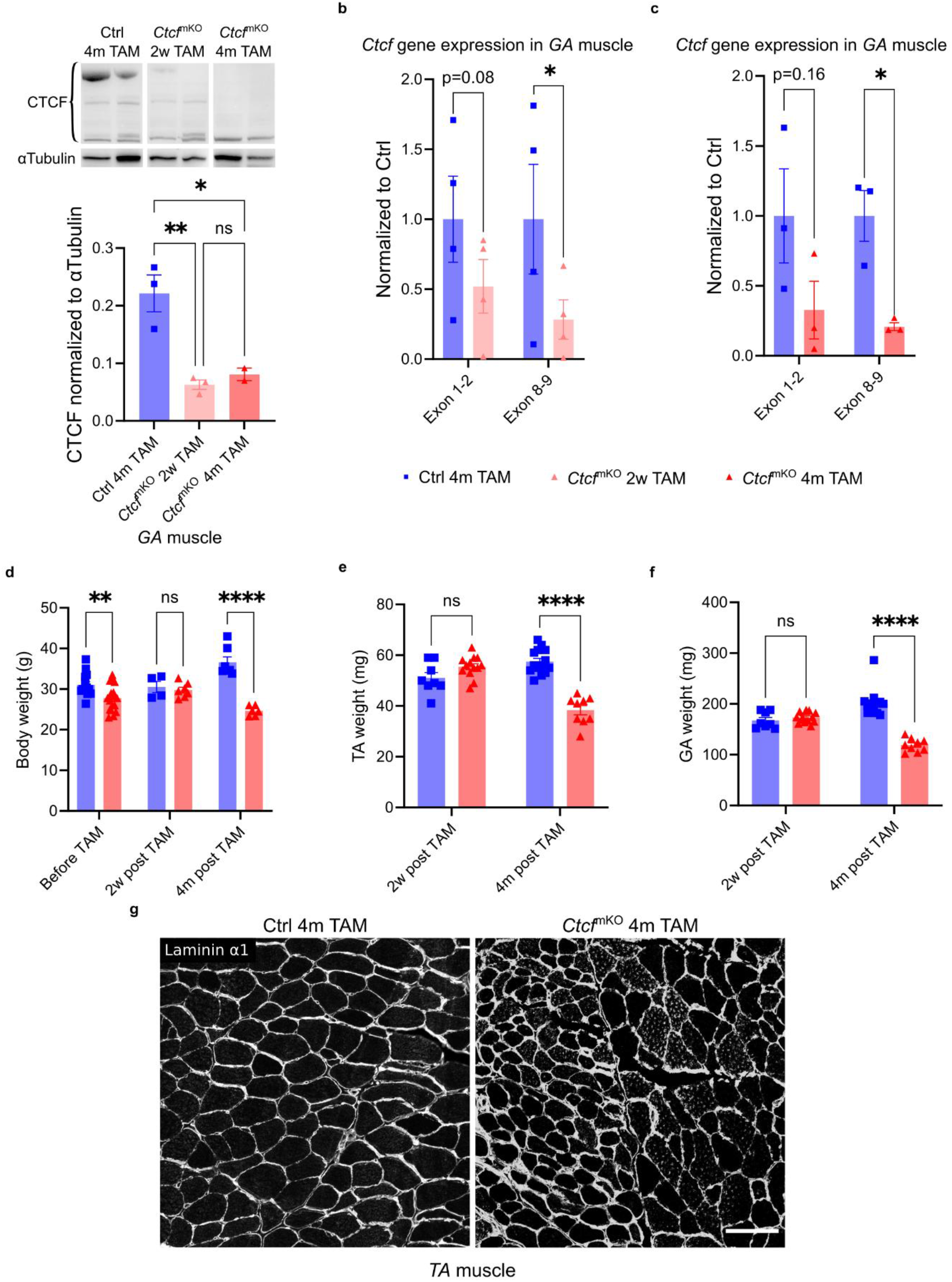
Adult myofiber-specific Ctcf depletion recapitulates the constitutive depletion phenotype. a, Western blot of CTCF protein in GA muscle of Ctrl 4 months tamoxifen (TAM; blue squares) and CtcfmKO (2 weeks TAM as pink triangles and 4 months TAM as red triangles) mice, western blot membrane (upper panel), quantification to αTubulin (lower panel). n = 3; bars represent mean±SEM; P values from two-tailed unpaired T test. b-c, RT-qPCR of Ctcf gene expression in whole GA muscle of Ctrl 4 months post tamoxifen (TAM; blue squares) and CtcfmKO mice 2 weeks post TAM as pink triangles (b),; or 4 months post TAM as red triangles (c). n = 5-8 mice; bars represent mean±SEM; P values from two-tailed unpaired T test. d, Body weight of Ctrl (blue squares) and CtcfmKO (red triangles) mice before, 2 weeks and 4 months post TAM. n = 4-18 mice; P values from two-tailed unpaired T test. e, Weight of the TA muscle of Ctrl (blue squares) and CtcfmKO (red triangles) mice at 2 weeks and 4 months post TAM. n = 8-14 muscles; P values from two-tailed unpaired T test. f, Weight of the GA muscle (top) and TA muscle (bottom) of Ctrl (blue squares) and CtcfmKO (red triangles) mice at 2 weeks and 4 months post TAM. n = 8-14 muscles; P values from two-tailed unpaired T test. g, Representative immunostaining of Laminin α1 in transversal section of the TA muscle of Ctrl (left) and CtcfmKO (right) at 4 months post TAM; scale bar = 100µm. *P<0.05; ** P<0.01; **** P<0.001; ns = non significant.

**Fig. S4.**
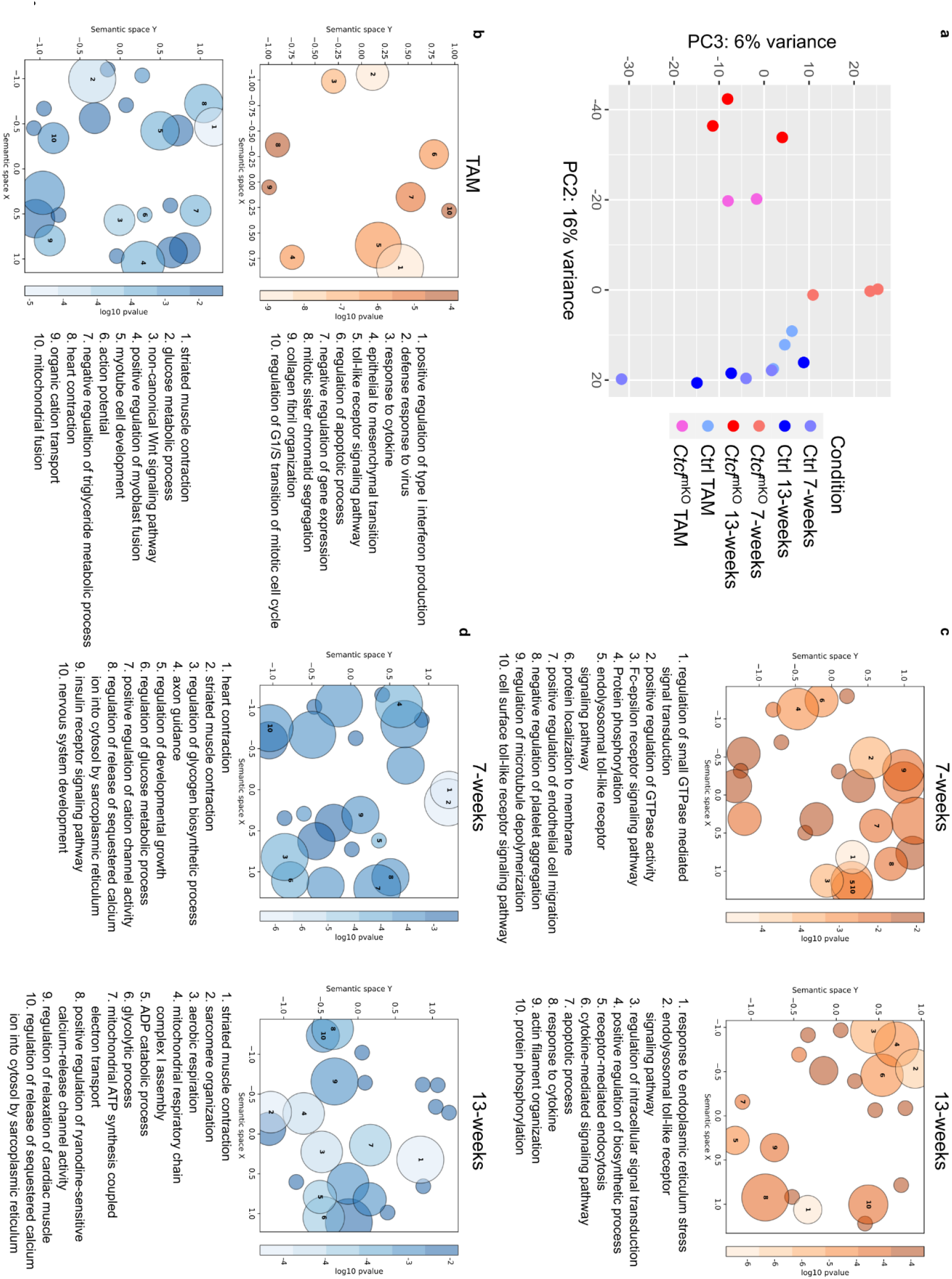
Muscle transcriptional changes are similar in constitutive and adult myofiber-specific Ctcf depletion. a, Principal Component Analysis (PCA; in the current plot, PC2 and PC3 components) plot of the bulk RNA-seq datasets from whole GA muscle for the constitutive (at 7 and 13 weeks) and conditional (TAM) Ctrl and CtcfmKO mouse model 4 months post tamoxifen. n = 2-3 mice. b, Top 10 biological processes linked to upregulated (top, orange) or downregulated (bottom, blue) genes, in conditional CtcfmKO vs Ctrl 4 months post TAM. Bubbles represent gene ontology terms aggregated by semantic similarity with the GOFigure python package. DEGs defined by |log2FC| > 0.25 and P value < 0.05. n = 2-3 mice. c, Top 10 biological processes linked to upregulated genes in constitutive CtcfmKO vs Ctrl at 7 (left) and 13 weeks (right). Bubbles represent gene ontology terms aggregated by semantic similarity with the GOFigure python package. DEGs defined by |log2FC| > 0.25 and P value < 0.05. n = 3 mice. d, Top 10 biological processes linked to downregulated genes in constitutive CtcfmKO vs Ctrl at 7 (left) and 13 weeks (right). Bubbles represent gene ontology terms aggregated by semantic similarity with the GOFigure python package. DEGs defined by |log2FC| > 0.25 and P value < 0.05. n = 2-3 mice.

**Fig. S5.**
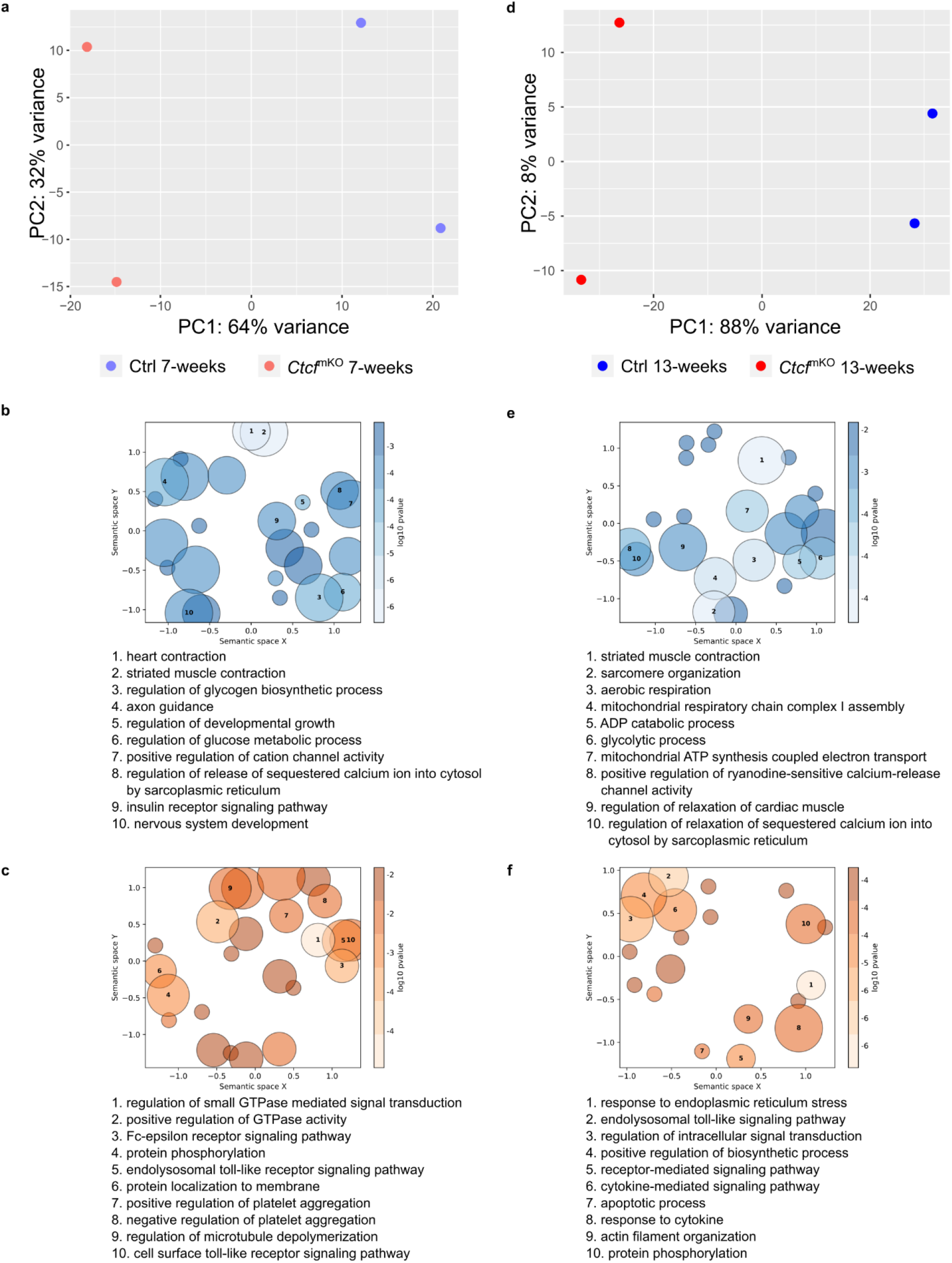
Pseudo-bulk differential analysis of all myonuclei. a,d, PCA plot of pseudo-bulk snRNA-seq from myonuclei at 7 (a) and 13 (d) weeks in CtcfmKO and Ctrl. n = 2. b-c, Top 10 biological processes linked to downregulated (b, blue) or upregulated (c, orange) genes in myonuclei at 7 weeks. Bubbles represent gene ontology terms aggregated by semantic similarity with the GOFigure python package. DEGs defined by |log2FC| > 0.25 and P value < 0.05. n = 2 mice. e-f, Top 10 biological processes linked to downregulated (e, blue) or upregulated (f, orange) genes in myonuclei at 13 weeks. Bubbles represent gene ontology terms aggregated by semantic similarity with the GOFigure python package. DEGs defined by |log2FC| > 0.25 and P value < 0.05. n = 2 mice.

**Fig. S6.**
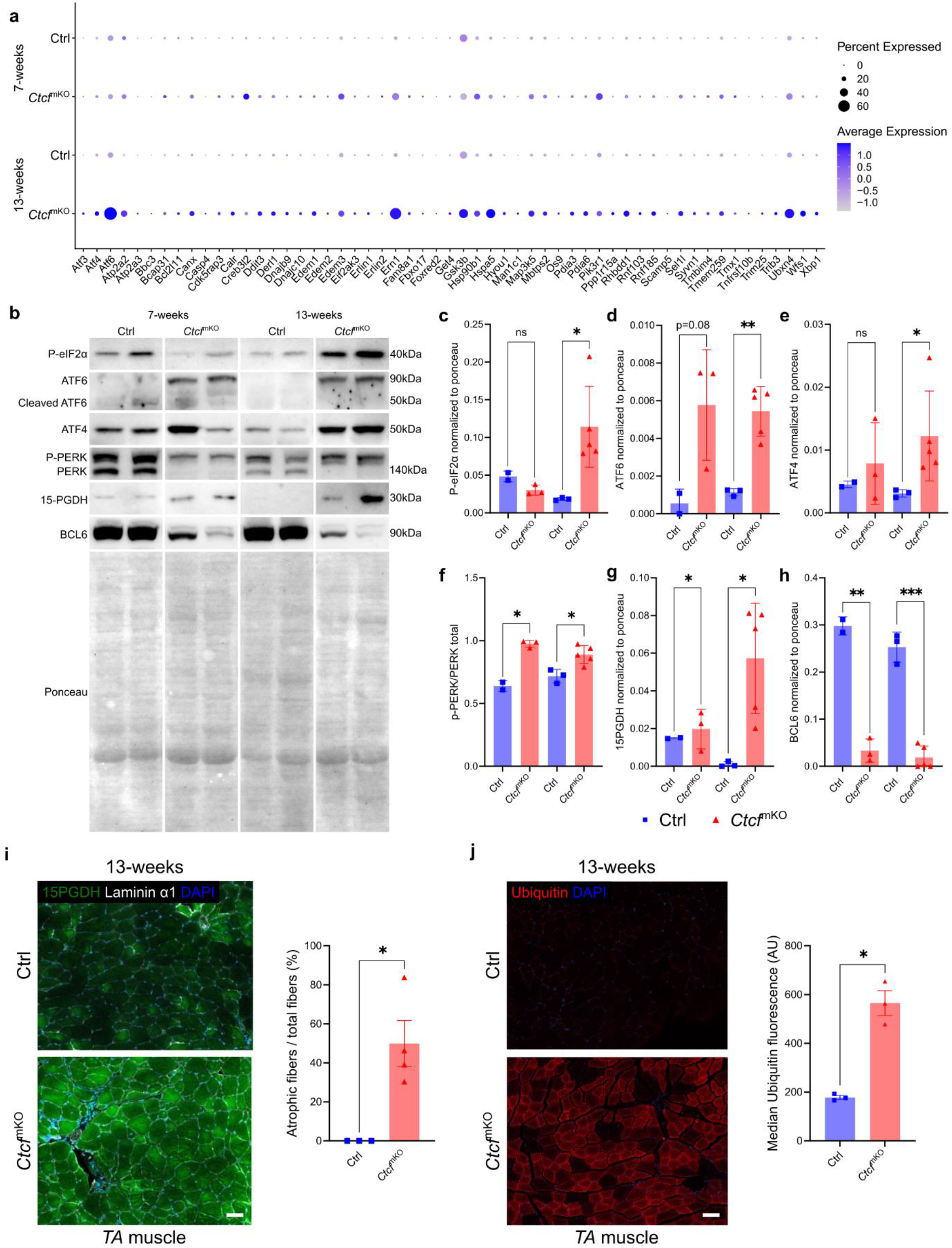
Ctcf depletion induces ER stress, unfolded protein response activation and proteostatic alterations in skeletal muscle. a, Dotplot analysis of ER/UPR signaling genes defined by gene ontology of biological processes in the myonuclei between Ctrl and CtcfmKO mice at 7 and 13 weeks. n = 2 mice. b, Western blot membranes of Ser51-phosphorylated eIF2α (P-eIF2α), ATF6, ATF4, Phosphorylated PERK (P-PERK), PERK, 15-PGDH, BCL6 and Ponceau control in GA muscle of Ctrl and CtcfmKO mice at 7 (left) and 13 weeks (right). c-h, Normalization of the western blot for Ser51-phosphorylated eIF2α (P-eIF2α), ATF6, ATF4, 15-PGDH and BCL6 over Ponceau in GA muscle of Ctrl (blue squares) and CtcfmKO (red triangles) mice at 13 weeks. Normalization of the phosphorylation of PERK is compared to the total-PERK in f. n = 2-5; bars are mean±SEM; P values of two-tailed unpaired T test. i, Representative images of CODEX of transversal sections of TA muscle of Ctrl and CtcfmKO at 13 weeks (left). 15-PGDH in green, Laminin α1 in white and DAPI in blue. Scale bar = 100µm. Analysis of the percentage of muscle fibers presenting 15-PGDH expression in TA muscle of Ctrl (blue squares) and CtcfmKO (red triangles) at 13 weeks (right). n = 3-4 mice; bars are mean±SEM; P values of two-tailed unpaired T test. j, Representative images of CODEX of transversal sections of TA muscle of Ctrl and CtcfmKO at 13 weeks (left). Ubiquitin in red and DAPI in blue. Scale bar = 100µm. Quantification of median of Ubiquitin in the TA muscle of Ctrl (blue squares) and CtcfmKO (red triangles) at 13 weeks (right). n = 3-4 mice; bars are mean±SEM; P values of two-tailed unpaired T test. *P<0.05; ** P<0.01; **** P<0.001; ns = non significant.

**Fig. S7.**
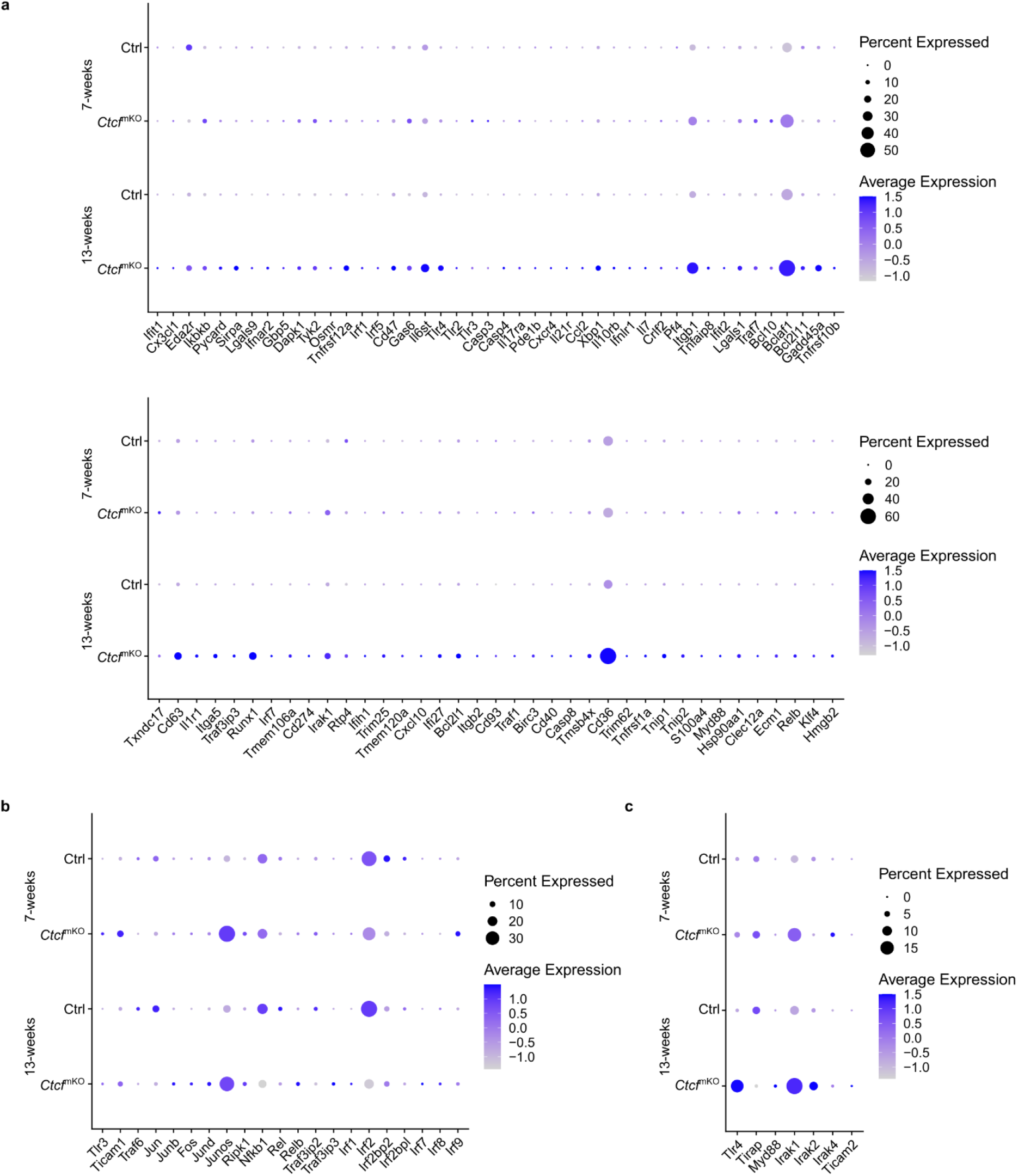
Ctcf depletion activates Toll-like receptor and NF-κB signaling in myonuclei. a, Dotplot analysis of immune signaling genes defined by gene ontology of biological processes in the myonuclei between Ctrl and CtcfmKO mice at 7 and 13 weeks. n = 2 mice. b, Dotplot analysis of Tlr3 signaling related gene expression in the myonuclei between Ctrl and CtcfmKO mice at 7 and 13 weeks. n = 2 mice. c, Dotplot analysis of Tlr4 signaling related gene expression in the myonuclei between Ctrl and CtcfmKO mice at 7 and 13 weeks. n = 2 mice.

**Fig. S8.**
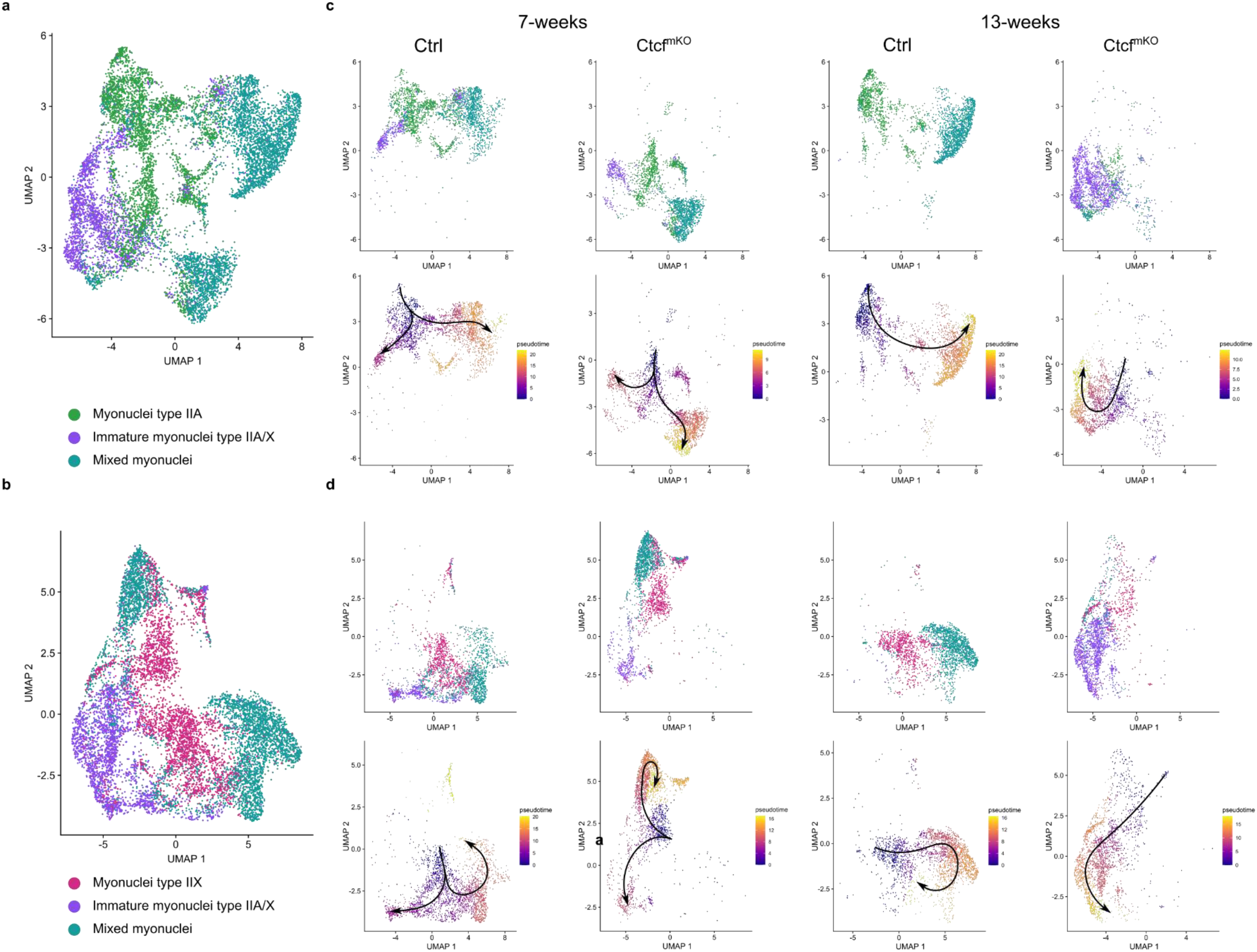
Ctcf depletion altered myonuclei type IIA and IIX pseudotime trajectories. a, UMAP of Type IIA, Immature type IIA/IIX and Mixed myonuclei clusters. n = 2 mice. b, UMAP of Type IIX, Immature type IIA/IIX and Mixed myonuclei clusters. n = 2 mice. c, UMAP of Type IIA, Immature type IIA/IIX and Mixed myonuclei clusters split by conditions (Ctrl and CtcfmKO at 7 and 13 weeks; top), colored and labeled by subclusters. Monocle3 pseudotime UMAP of Type IIA, Immature type IIA/IIX and Mixed myonuclei clusters split by conditions (Ctrl and CtcfmKO at 7 and 13 weeks; bottom). n = 2 mice. d, UMAP of Type IIX, Immature type IIA/IIX and Mixed myonuclei clusters split by conditions (Ctrl and CtcfmKO at 7 and 13 weeks; top), colored and labeled by subclusters. Monocle3 pseudotime UMAP of Type IIX, Immature type IIA/IIX and Mixed myonuclei clusters split by conditions (Ctrl and CtcfmKO at 7 and 13 weeks; bottom). n = 2 mice.

**Fig. S9.**
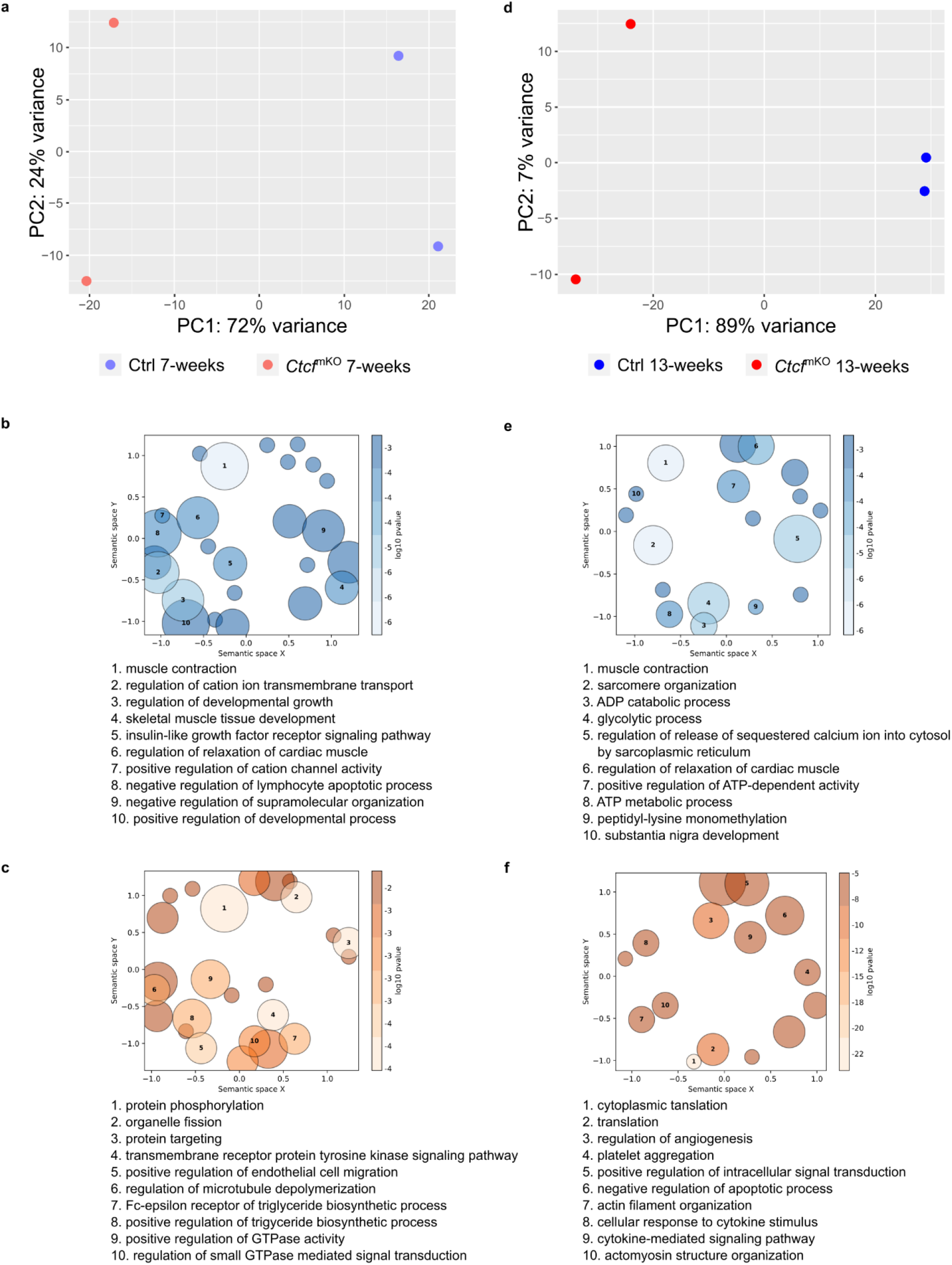
Pseudo-bulk differential analysis of myonuclei type IIB. a,d, PCA plot of pseudo-bulk RNA-seq from myonuclei type IIB at 7 (a) and 13 (d) weeks in CtcfmKO and Ctrl. n = 2 mice. b-c, Top 10 biological processes linked to downregulated (b, blue) or upregulated (c, orange) genes in myonuclei at 7 weeks. Bubbles represent gene ontology terms aggregated by semantic similarity with the GOFigure python package. DEGs defined by |log2FC| > 0.25 and P value < 0.05. n = 2 mice. e-f, Top 10 biological processes linked to downregulated (e, blue) or upregulated (f, orange) genes in myonuclei at 13 weeks. Bubbles represent gene ontology terms aggregated by semantic similarity with the GOFigure python package. DEGs defined by |log2FC| > 0.25 and P value < 0.05. n = 2 mice.

**Fig. S10.**
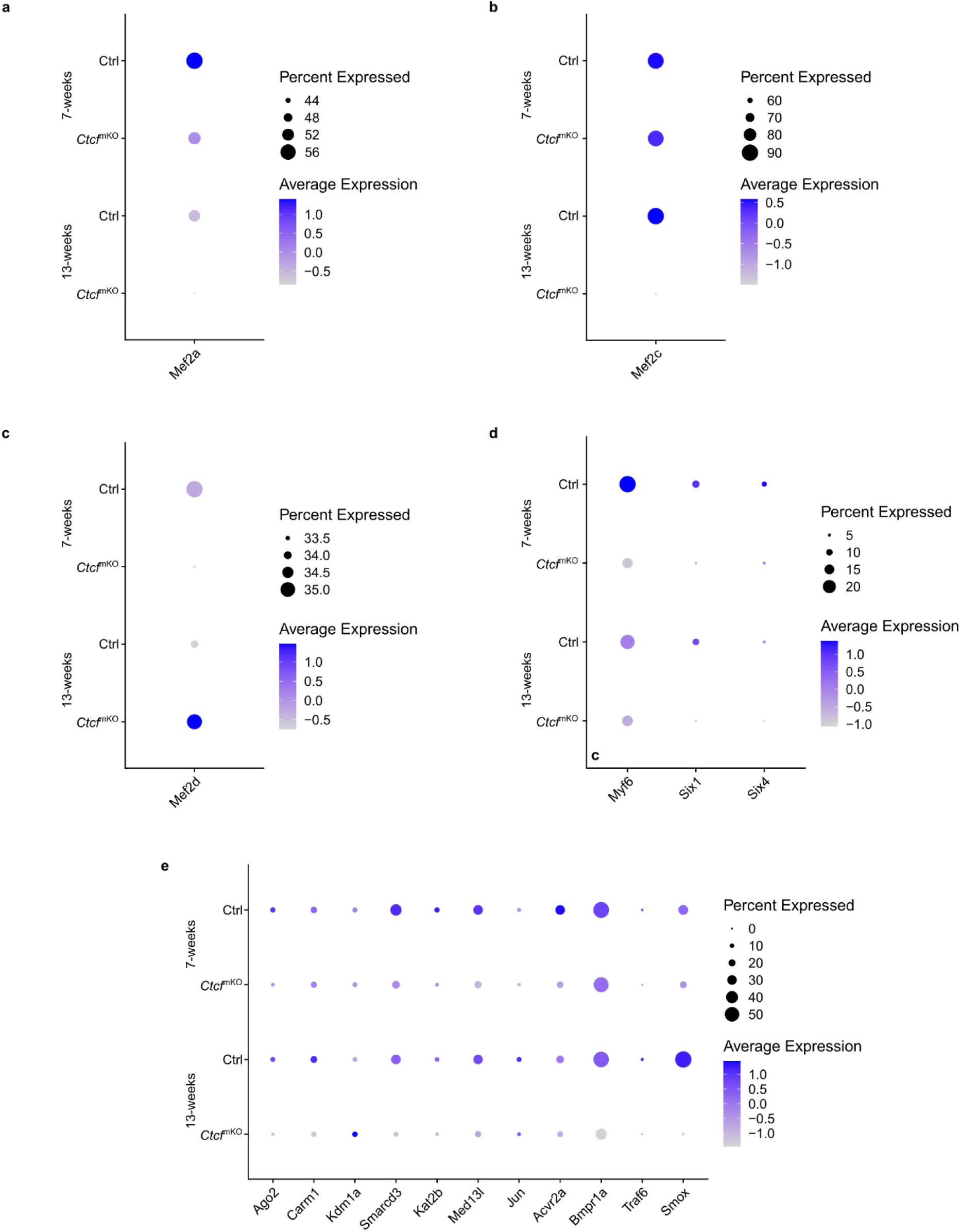
Muscle transcription factors expression is altered by Ctcf depletion in myonuclei type IIB. a, Dotplot analysis of Mef2a gene in myonuclei type IIB between Ctrl and CtcfmKO mice at 7 and 13 weeks. n = 2 mice. b, Dotplot analysis of Mef2c gene in myonuclei type IIB between Ctrl and CtcfmKO mice at 7 and 13 weeks. n = 2 mice. c, Dotplot analysis of Mef2d gene in myonuclei type IIB between Ctrl and CtcfmKO mice at 7 and 13 weeks. n = 2 mice. d, Dotplot analysis of Myf6, Six1 and Six4 genes in myonuclei type IIB between Ctrl and CtcfmKO mice at 7 and 13 weeks. n = 2 mice. e, Dotplot analysis of transcription factors and muscle co-activator genes in myonuclei type IIB between Ctrl and CtcfmKO mice at 7 and 13 weeks. n = 2 mice.

**Fig. S11.**
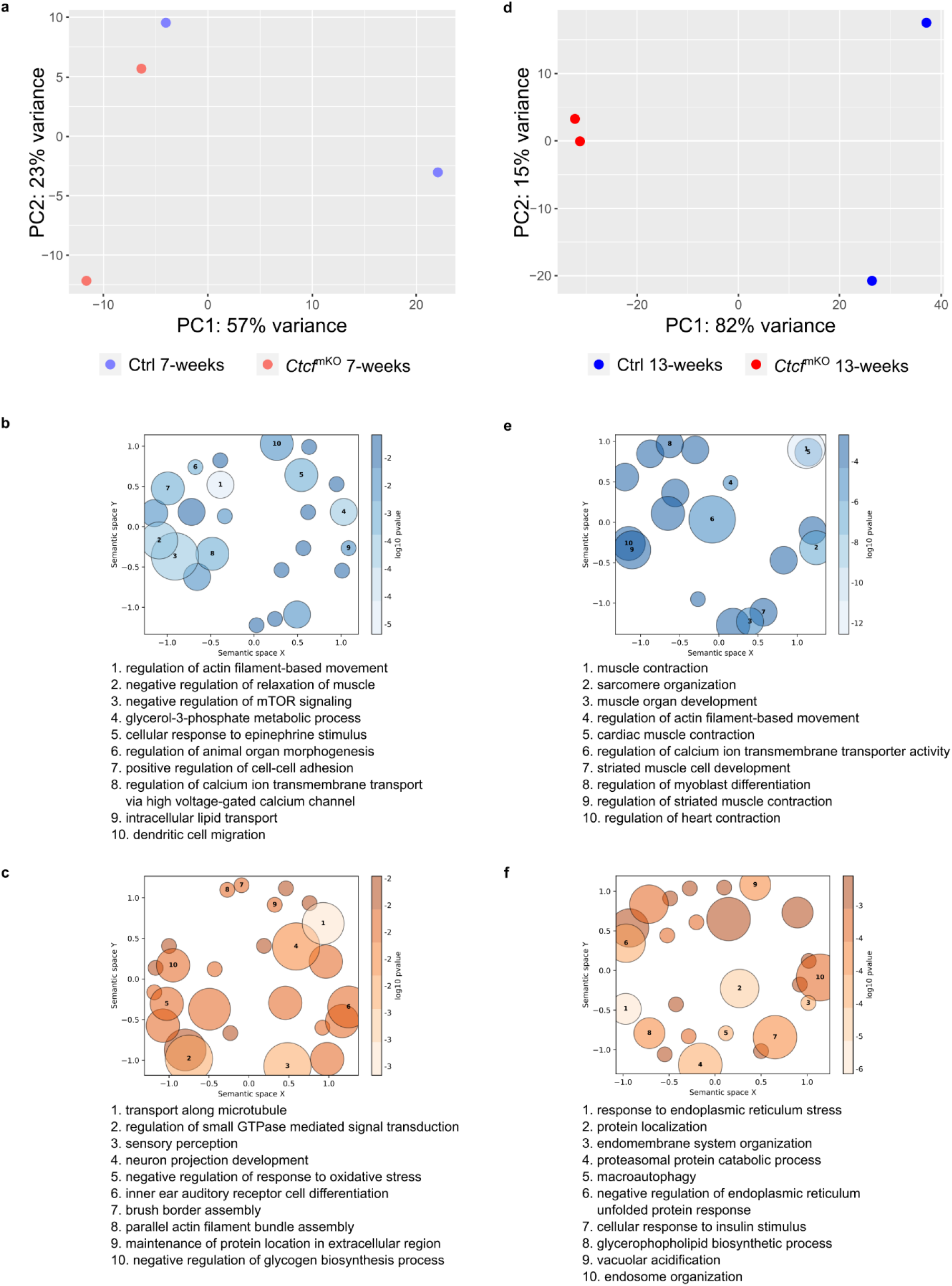
Pseudo-bulk differential analysis of “other myonuclei”. a,d, PCA plot of pseudo-bulk RNA-seq from the “other myonuclei” at 7 (a) and 13 (d) weeks in CtcfmKO and Ctrl. n = 2 mice. b-c, Top 10 biological processes linked to downregulated (b, blue) or upregulated (c, orange) genes in myonuclei at 7 weeks. Bubbles represent gene ontology terms aggregated by semantic similarity with the GOFigure python package. DEGs defined by |log2FC| > 0.25 and P value < 0.05. n = 2 mice. e-f, Top 10 biological processes linked to downregulated (e, blue) or upregulated (f, orange) genes in myonuclei at 13 weeks. Bubbles represent gene ontology terms aggregated by semantic similarity with the GOFigure python package. DEGs were defined as genes with |log2FC|>0.25 and P value < 0.05. n = 2 mice.

**Fig. S12.**
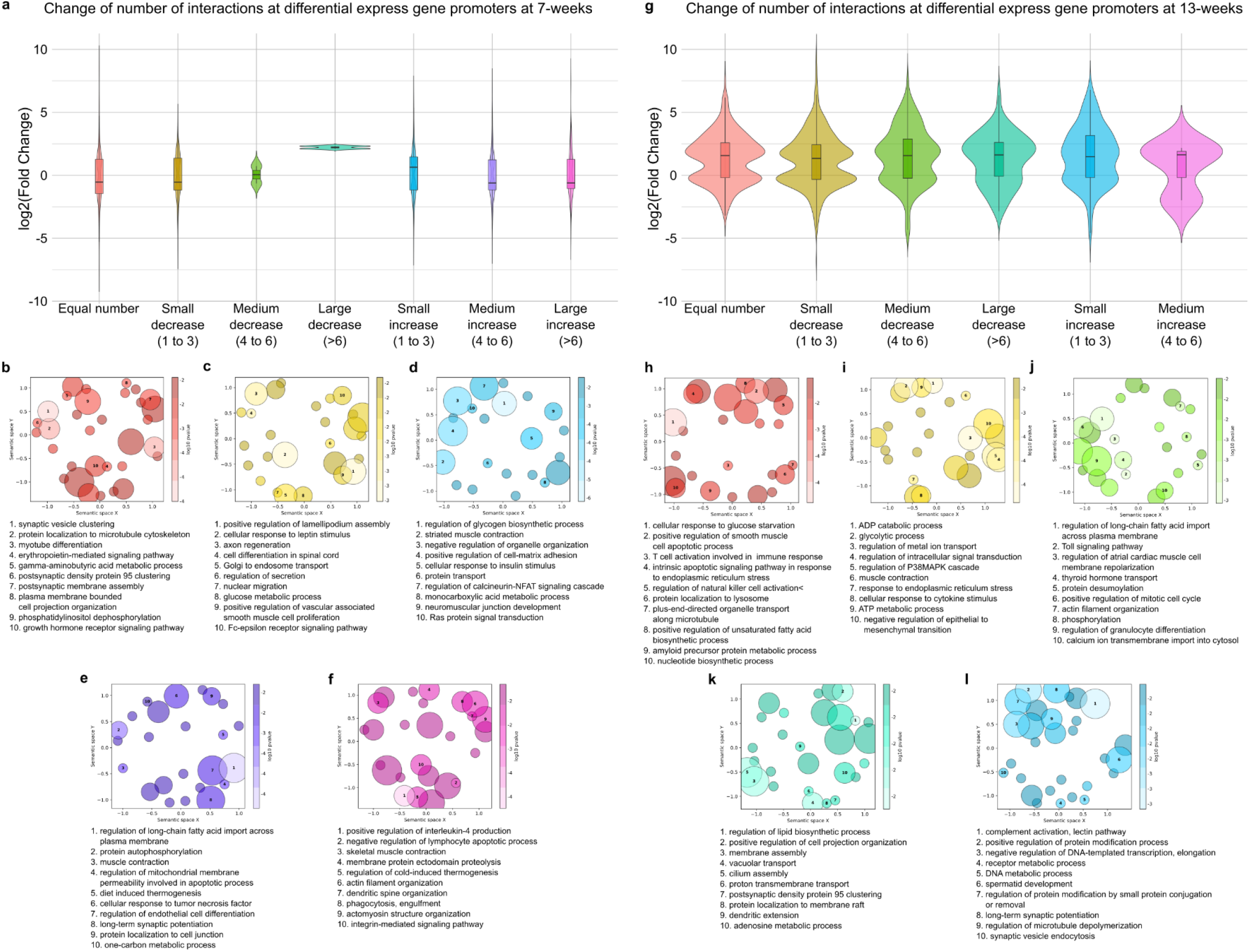
Ctcf loss disrupts the coupling between chromatin interactions and transcription at 7 and 13 weeks. a, Violin plot of the log2fold change distribution for differentially expressed genes that change number of interaction partners (classified as small, medium and large decrease or increase) or not (equal number) upon CTCF loss at 7 weeks. b-f, Top 10 biological processes linked to the classes represented in a. Specifically, b shows biological processes of DEGs with Equal number of interactions, c, biological processes of DEGs showing a small decrease in interaction partners, or a small increase in d, medium increase in e, and large increase in f. Bubbles represent gene ontology terms aggregated by semantic similarity with the GOFigure python package. g, Violin plot of the log2fold change distribution for differentially expressed genes that change number of interaction partners (classified as small, medium or large decrease or increase) or not (equal number) upon CTCF loss at 13 weeks. h-l, Top 10 biological processes linked to the classes represented in g. Specifically, h shows biological processes of DEGs with Equal number of interactions, i, biological processes of DEGs showing a small decrease in interaction partners, medium decrease in j, large decrease in k and small increase in l. Bubbles represent gene ontology terms aggregated by semantic similarity with the GOFigure python package.

**Fig. S13.**
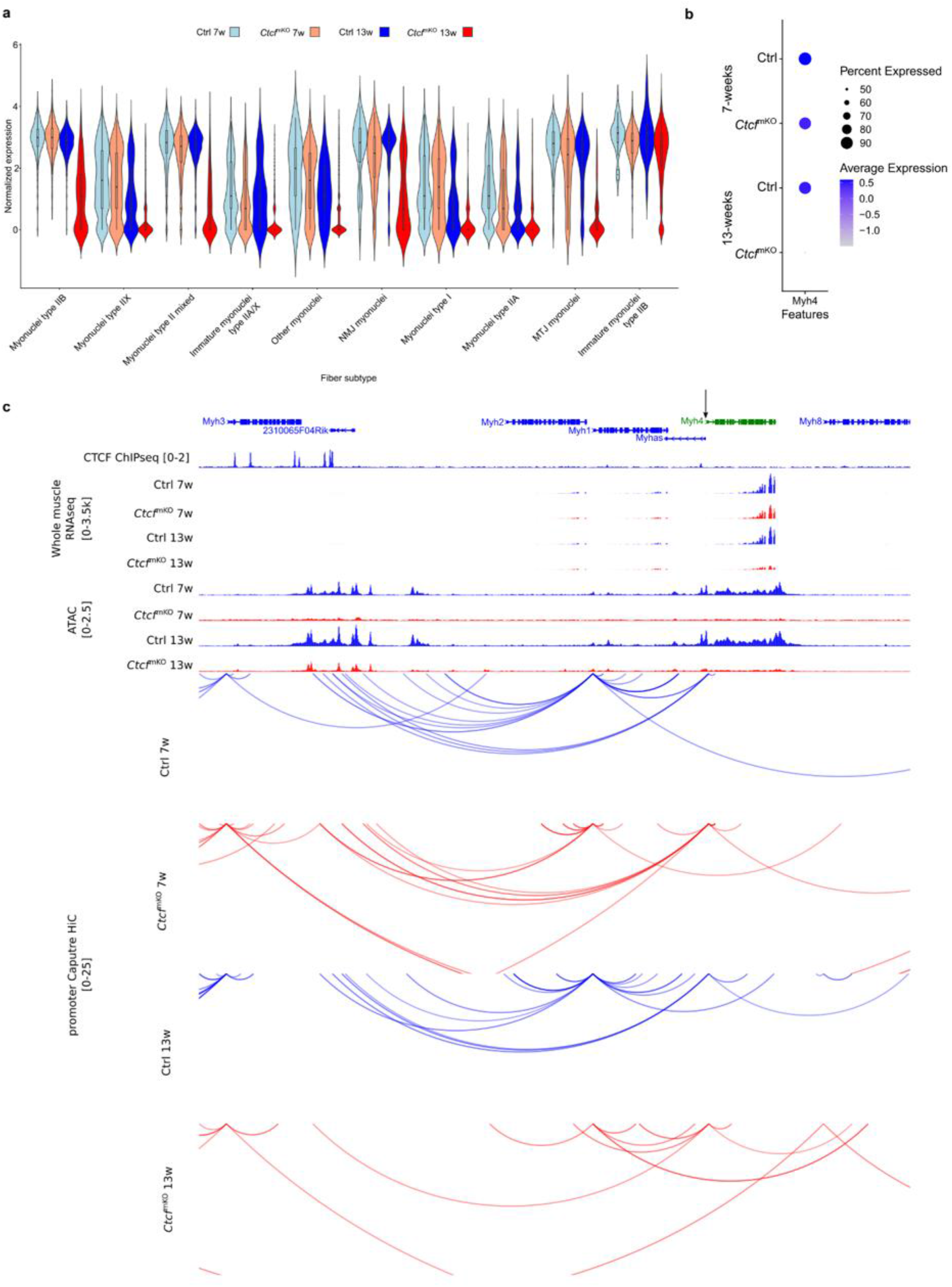
Ctcf loss induces chromatin rewiring at the Myh4 locus. a, Violin plot of the normalized gene expression of Myh4 in the different snRNAseq subcluster of the myonuclei metacluster between Ctrl and CtcfmKO mice at 7 and 13 weeks. n = 2 mice. b, Dotplot analysis of Myh4 in the myonuclei between Ctrl and CtcfmKO mice at 7 and 13 weeks. c, WashU screenshot of the genomic regions of Myh4. Gene of interest (GOI) is shown in green, GOI promoter is highlighted by an arrow. Tracks from top to bottom: refseq gene, RNA-seq tracks in Ctrl (blue) and CtcfmKO (red) muscles at 7 and 13 weeks, ATAC-seq tracks in Ctrl myonuclei (blue) and CtcfmKO (red) at 7 and 13 weeks, CTCF ChIP-seq in isolated myonuclei from WT mice (black), promoter capture HiC interaction in Ctrl myonuclei (blue) and CtcfmKO (red) at 7 and 13 weeks.

**Table S1.**
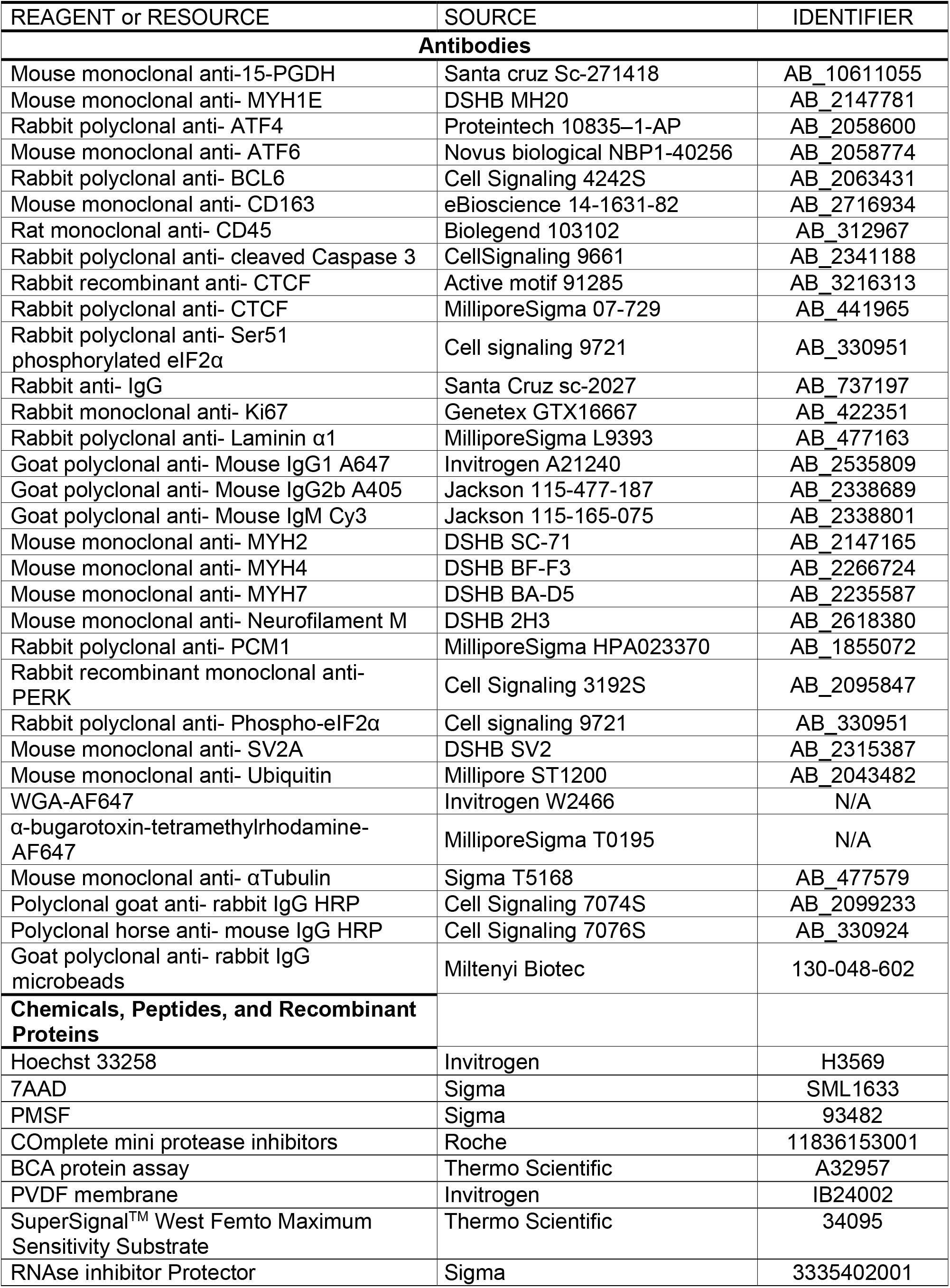
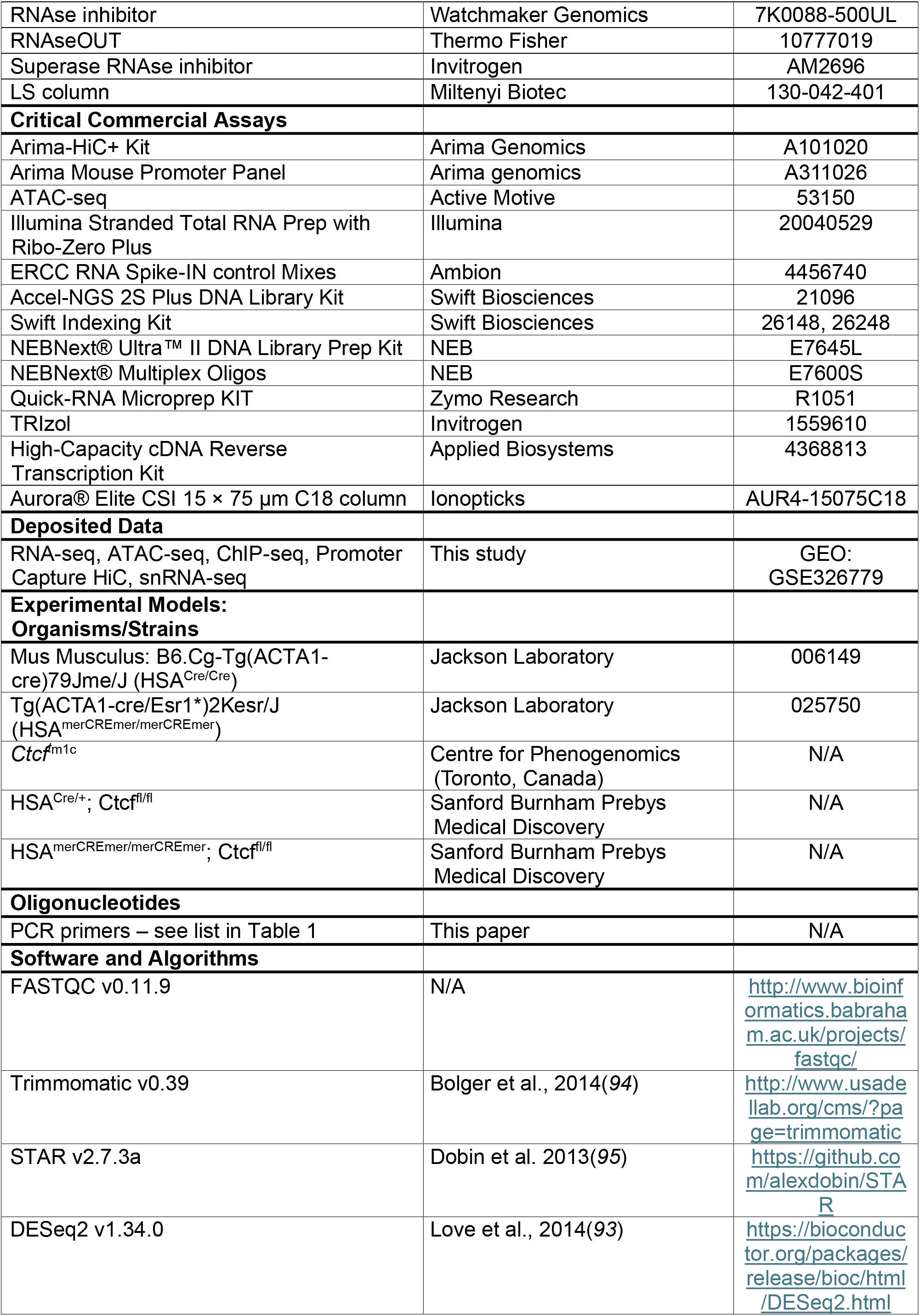
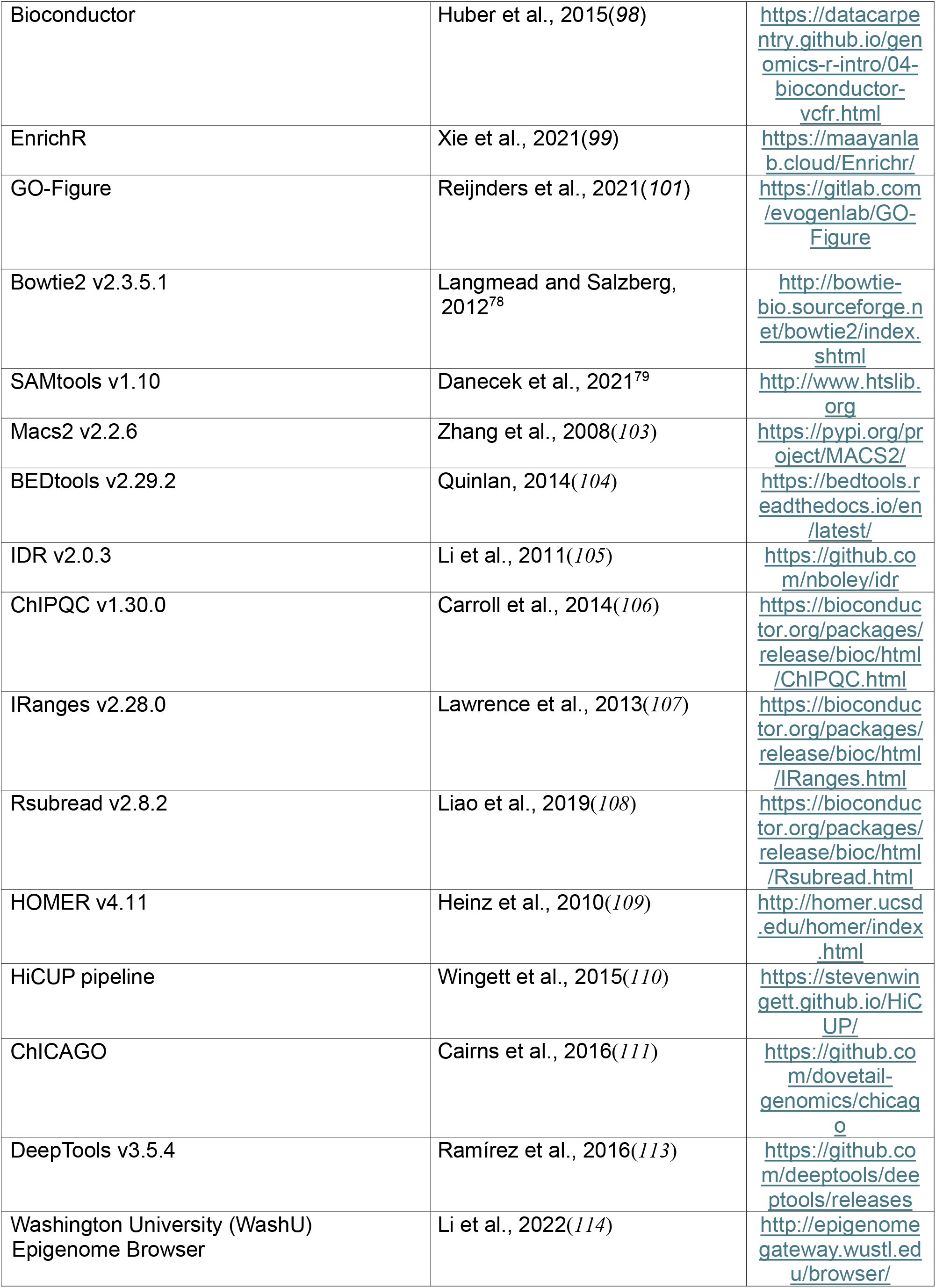
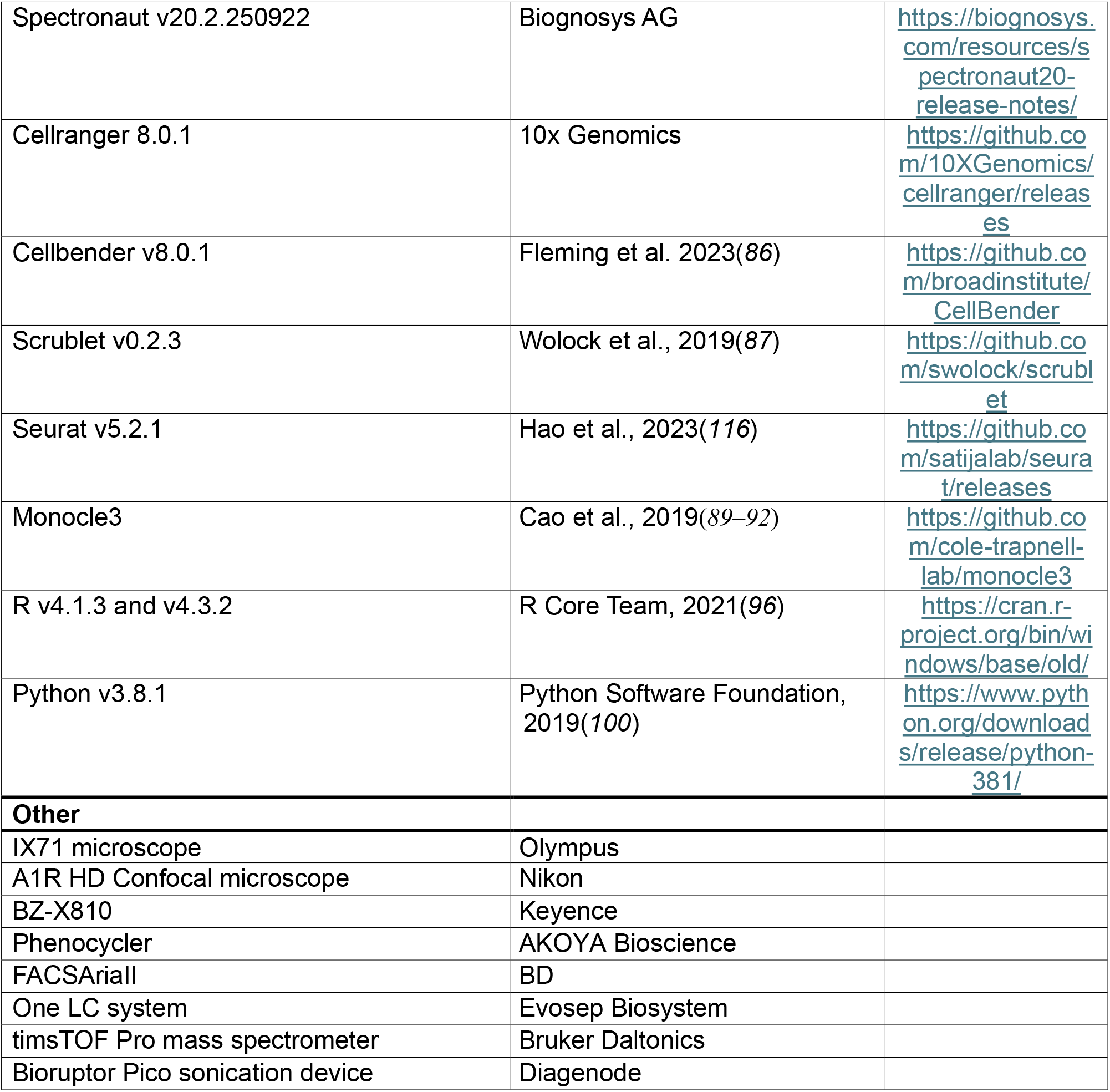
Key resources table.

